# Inflammatory proteolysis generates pathogenic APOL1 fragments with distinct intracellular toxicities in podocytes derived from children with HIV associated nephropathy

**DOI:** 10.64898/2026.08.12.744497

**Authors:** Jinliang Li, Jing Yu, Jharna Das, Lian Xu, Pankaj Kumar, Zhe Han, Patricio E. Ray

## Abstract

APOL1 risk variants are the strongest genetic determinants of HIV-associated nephropathy (HIVAN), yet the mechanisms linking inflammation to APOL1-mediated podocyte injury remain poorly understood because authentic patient-derived human disease models are lacking. Using urine-derived podocytes established from children with HIVAN and endogenous APOL1 reporter cell lines derived from these cells, we identified a previously unrecognized pathway of inflammatory, cathepsin-dependent APOL1 proteolysis. Endogenous APOL1 cleavage was detected in patient-derived podocytes, whereas reporter cell lines enabled the identification and functional characterization of N-terminal and C-terminal APOL1 fragments with distinct intracellular localization and pathogenic functions. The nuclear N-terminal fragment activated inflammatory transcriptional programs and promoted podocyte injury, whereas the membrane-associated C-terminal fragment mediated membrane toxicity and remained susceptible to pharmacologic inhibition by inaxaplin. Cathepsin S directly cleaved APOL1 in vitro, linking inflammatory signaling to APOL1 fragmentation. These findings identify inflammatory APOL1 proteolysis as a mechanism that partitions APOL1 toxicity into distinct pathogenic programs and nominate APOL1 processing as a therapeutic target for HIV-associated and other APOL1-mediated kidney diseases.

## Introduction

APOL1 risk variants are among the strongest inherited determinants of kidney disease identified in humans. ^1–3^ Individuals carrying two APOL1 risk alleles have markedly increased susceptibility to focal segmental glomerulosclerosis (FSGS), HIV-associated nephropathy (HIVAN), collapsing glomerulopathy, and other progressive kidney diseases ^1–3^ The G1 and G2 variants account for a substantial proportion of kidney disease among individuals of West African ancestry and have transformed our understanding of genetic susceptibility to chronic kidney disease. ^4,5^ Despite compelling genetic evidence, the mechanisms by which APOL1 drives podocyte injury and progressive kidney disease remain incompletely understood, particularly in childhood HIV-associated nephropathy (HIVAN).

Current models of APOL1 toxicity focus primarily on the full-length protein. APOL1 localizes to multiple intracellular membranes and has been linked to ion transport, membrane permeabilization, vesicular trafficking defects, mitochondrial dysfunction, and activation of cellular stress pathways. ^6–19^ These observations support a prevailing model in which disease is driven largely by aberrant activities of intact APOL1. ^20^ However, this framework does not readily explain the context dependence of APOL1 in HIVAN, the selective vulnerability of podocytes, or the diversity of cellular phenotypes associated with APOL1 expression. Although interactions with APOL3 may also contribute to toxicity,^10,13^ the assumption that full-length APOL1 is the principal pathogenic species remains largely untested.

Inflammation is a central feature of APOL1-associated kidney disease. Interferons are potent inducers of APOL1 expression and have been implicated in HIVAN, lupus nephritis, virus-associated collapsing glomerulopathy, COVID-19–associated nephropathy, and related disorders. ^14,21–24^ While inflammatory activation is widely viewed as a key determinant of APOL1-mediated injury, whether it alters APOL1 through regulated post-translational mechanisms remains unknown.

Proteolytic processing is a fundamental mechanism by which proteins acquire new functions, localization patterns, and pathogenic properties.^25^ Despite extensive investigation of APOL1 trafficking, membrane biology, channel activity, and interactions with APOL3, whether APOL1 undergoes regulated proteolysis and whether APOL1-derived fragments contribute to disease pathogenesis have remained largely unexplored.

Addressing these questions is challenging because APOL1 is a relatively recent primate gene absent from rodents.^26^ HIV-associated nephropathy (HIVAN) provides a particularly informative context because it is strongly APOL1-dependent and characterized by chronic interferon activation, robust APOL1 induction, and severe podocyte injury.^3^ Pediatric HIVAN also provides access to disease-relevant human podocytes while minimizing confounding age-related comorbidities and secondary renal injury.^27^ To address these challenges, we leveraged a unique human experimental platform comprising patient-derived podocytes established from children with HIVAN and the first endogenous CRISPR-engineered GFP-APOL1 reporter derivatives generated from these cells.

Here, we integrate analyses of pediatric HIVAN kidney biopsies, urine-derived primary podocytes, endogenous CRISPR-engineered APOL1 reporter cell lines, transcriptomics, and biochemical approaches to define APOL1 biology in disease-relevant human cells. We identify an inflammatory, cathepsin-dependent pathway of APOL1 proteolysis that generates N-terminal and C-terminal fragments with distinct localization, biological activities, and pharmacologic responses. APOL1 proteolysis generates pathogenic nuclear APOL1 species that activate inflammatory transcriptional programs and promote podocyte injury.

APOL1 processing is associated with the emergence of pathogenic nuclear APOL1 species, activation of inflammatory transcriptional programs, and enhanced podocyte injury. Finally, the APOL1 inhibitor inaxaplin selectively modulates specific APOL1 fragments, suggesting that therapeutic responses may depend on APOL1 fragment composition in addition to full-length APOL1 abundance. Together, these findings identify inflammatory APOL1 proteolysis as a previously unrecognized mechanism that partitions APOL1 toxicity into distinct intracellular pathogenic programs.”

## Results

### APOL1 is expressed in podocytes from children with HIVAN and is induced by inflammatory cytokines

To define the localization of APOL1 in pediatric HIV-associated nephropathy (HIVAN), we analyzed kidney sections from children with HIVAN and HIV-infected controls. Immunohistochemical staining demonstrated prominent APOL1 expression in podocytes, glomerular epithelial cells and endothelial cells in children living with HIV-1 (Fig. 1a-b). Renal sections from children without kidney disease exhibited stronger and more preserved glomerular endothelial APOL1 staining compared with those with HIVAN (Fig. 1a-b; n = 4 per group). Consistent with renal APOL1 expression, APOL1 was detected in urine samples obtained from children living with HIV. Urinary APOL1 concentrations were significantly elevated in those with HIV-chronic kidney disease (HIV-CKD) compared with those without kidney disease (HIV-C) (Fig. 1c).

**Figure 1.**
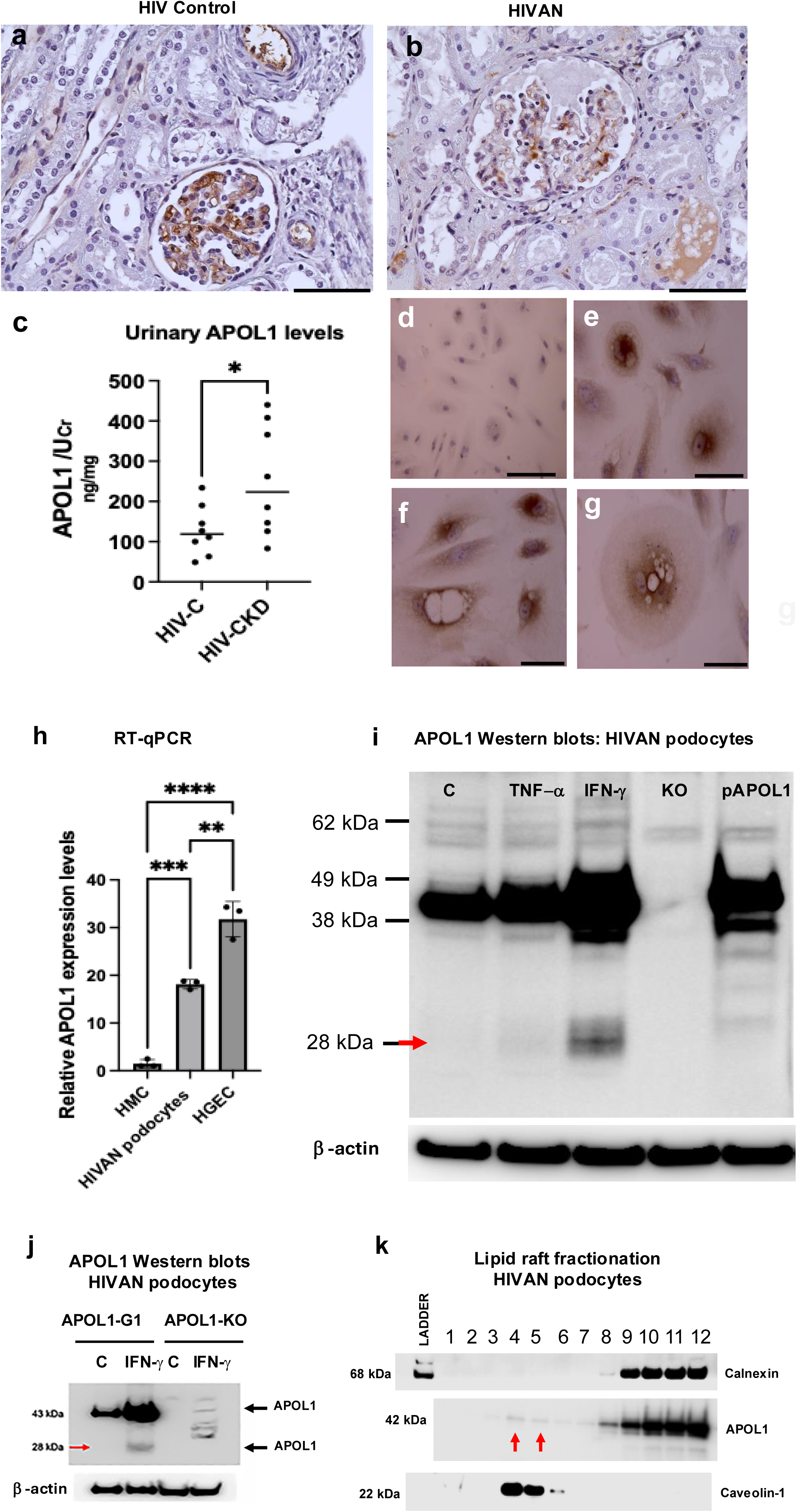
APOL1 expression in children with HIV-associated nephropathy and proteolytic processing of APOL1 in urine-derived HIVAN podocytes. **(a-b)** Representative APOL1 immunohistochemistry (IHC) in kidney sections from HIV-controls and HIVAN. APOL1 staining is detected mainly within glomerular cells and podocytes. Results are representative of four independent kidney sections per group. Scale bars, 50 μm, (c) Quantification of urinary APOL1 normalized to urinary creatinine (UCr) in HIV-control (HIV-C) and HIV-associated chronic kidney disease (HIV-CKD) subjects. Each dot represents an individual participant (*n* = 7 per group). P = 0.043. Student’s *t* test. (**d-g)** Representative APOL1 immunohistochemistry images of primary podocytes cultured from the urine of a child with HIVAN (HIVAN podocytes), showing heterogeneous cellular staining patterns. Panel d shows podocytes exposed to the control antibody. Panels e-g show podocytes exposed to the Sigma APOL1 antibody. Scale bars, 30 μm. **(h)** Relative APOL1 mRNA expression measured by RT-qPCR in human mesangial cells (HMCs), HIVAN podocytes, and human glomerular endothelial cells (HGECs). Data are representative of 3 independent biological samples per group. Mean ± SD (n= 3 per group). The expression of APOL1 mRNA in HIVAN podocytes was compared with HMC and HGEC by different cell types by one ANOVA with Turkey’s multiple comparisons ** P < 0.01; *** p< 0.001 **** P, 0.0001. **(i)** Immunoblot analysis of APOL1 in HIVAN podocytes treatment with TNF-α, IFN-γ, and homozygous APOL1-KO HIVAN podocytes transfected with an APOL1-G1 plasmid. β-actin served as a loading control. The red arrow indicates a lower-molecular-weight APOL1 fragment. Results are representative of 3 biological independent replicates per group. **(j)** Western blots showing APOL1-G1 expression in HIVAN podocytes and APOL1-KO HIVAN podocytes under basal conditions, and after IFN-γ stimulation. Full-length and a lower-molecular-weight APOL1 fragment (red arrow) are detected in HIVAN podocytes treated with IFN-γ but not detected in the APOL1-KO HIVAN podocytes treated in a similar manner. β-actin served as a loading control. Results are representative of two independent experiments. **(k)** Lipid raft fractionation of HIVAN podocytes showing distribution of APOL1 across sucrose-gradient fractions. Caveolin-1 and calnexin were used as markers of raft and non-raft fractions, respectively. Arrows indicate APOL1 detected within low-density membrane fractions. Results are representative of two independent biological replicates per group.

Podocytes cultured from the urine of children with HIVAN exhibited predominantly perinuclear APOL1 staining, although occasional nuclear localization was observed (Fig.1d-g; Supplementary Fig. 1 g-h). Primary HIVAN podocytes were shed in the urine in an undifferentiated state. Some clones rapidly differentiated in culture (Supplementary Fig 1 a, b, e, f, i, j), while others developed prominent vacuolization associated with marked perivacuolar and perinuclear APOL1 staining (Fig. 1 f-g; Supplementary Fig. 1 g-h). Unlike podocytes that are transfected with APOL1-G1, which died rapidly through cell swelling and detachment (Supplementary Fig. 1d), primary urinary HIVAN podocytes remained viable for a few days despite vacuolization and APOL1 redistribution, subsequently undergoing senescence and cell death (Fig.1 f-g; Supplementary Figs. 1c, g-h).

RT-qPCR studies detected APOL1 mRNA predominately in HIVAN podocytes and glomerular endothelial cells (HGEC), while almost undetectable levels were noted in human mesangial cells (HMC) (Fig 1h). Western blot analysis confirmed the expression of APOL1 protein in urinary HIVAN podocytes using specific antibodies from Genentech^28,29^, whereas APOL1-knockout (APOL1-KO) HIVAN podocytes showed no expression of APOL1 and therefore served as negative controls (Fig 1 i-j). IFN-γ induced a robust increase in APOL1 expression, while TNF-α produced a modest effect (Fig. 1i). Notably, IFN-γ and TNF-α stimulated HIVAN podocytes showed a lower-molecular-weight APOL1 fragment of approximately 28 kDa that was not detected under baseline conditions (control cells) or in APOL1-KO HIVAN podocytes under baseline conditions or treated with INF-γ (red arrows, Figs 1 i-j), suggesting cytokine-induced proteolytic processing of APOL1.

### APOL1-G1 localizes to lipid rafts

Because APOL1 exhibits high affinity for anionic phospholipids enriched in lipid rafts,^30^ we examined its membrane localization in parental HIVAN podocytes carrying the APOL1 G1/G1 genotype. Sucrose-gradient fractionation revealed endogenous APOL1-G1 signaling predominantly in the higher-density, non-raft fractions (fractions 8–12) in association with the calnexin ER marker, with limited signal in lipid raft fractions 4 and 5 (Fig 1k, red arrows). Caveolin-1, a lipid raft marker, was enriched in fractions 4-5 confirming successful isolation of detergent-resistant membrane domains (Fig. 1k). Additional studies done in HIVAN podocytes transiently transfected with a APOL1-G1 plasmid also confirmed the expression of APOL1-G1 in the same locations (Supplementary Fig. 3).

### Endogenous GFP-APOL1-G1 reporter HIVAN podocytes reveal distinct cytoplasmic and nuclear APOL1 states

To directly visualize APOL1 expressed from its endogenous locus, we generated GFP-tagged APOL1-G1 HIVAN podocyte lines using CRISPR-Cas9 genome editing (Fig. 2a). This approach avoids reliance on antibody-based detection methods, which have yielded divergent localization patterns in prior studies, including reports of intranuclear APOL1 localization^31^, and whose specificity has subsequently been questioned using APOL1-deficient cells.^28^ Endogenous tagging therefore enables direct assessment of APOL1 subcellular distribution at native expression levels while minimizing potential detection artifacts. Correct insertion of the GFP tag was confirmed by genomic sequencing, immunoblotting, and fluorescence microscopy. This strategy generated a GFP-APOL1-G1 fusion protein of approximately 70 kDa whose expression was confirmed by RNA-seq, RT-qPCR and immunoblotting (Fig. 2b-d). To assess potential off-target integration of the GFP donor cassette, whole-genome sequencing reads were analyzed for GFP-containing read pairs. All GFP-associated reads mapped to the APOL1 locus on chromosome 22, consistent with the intended insertion site. No evidence of GFP integration at other genomic locations was detected, supporting site-specific targeting of the APOL1 locus (Supplementary Tables. 1-2 and Supplementary Fig 2).

**Figure 2.**
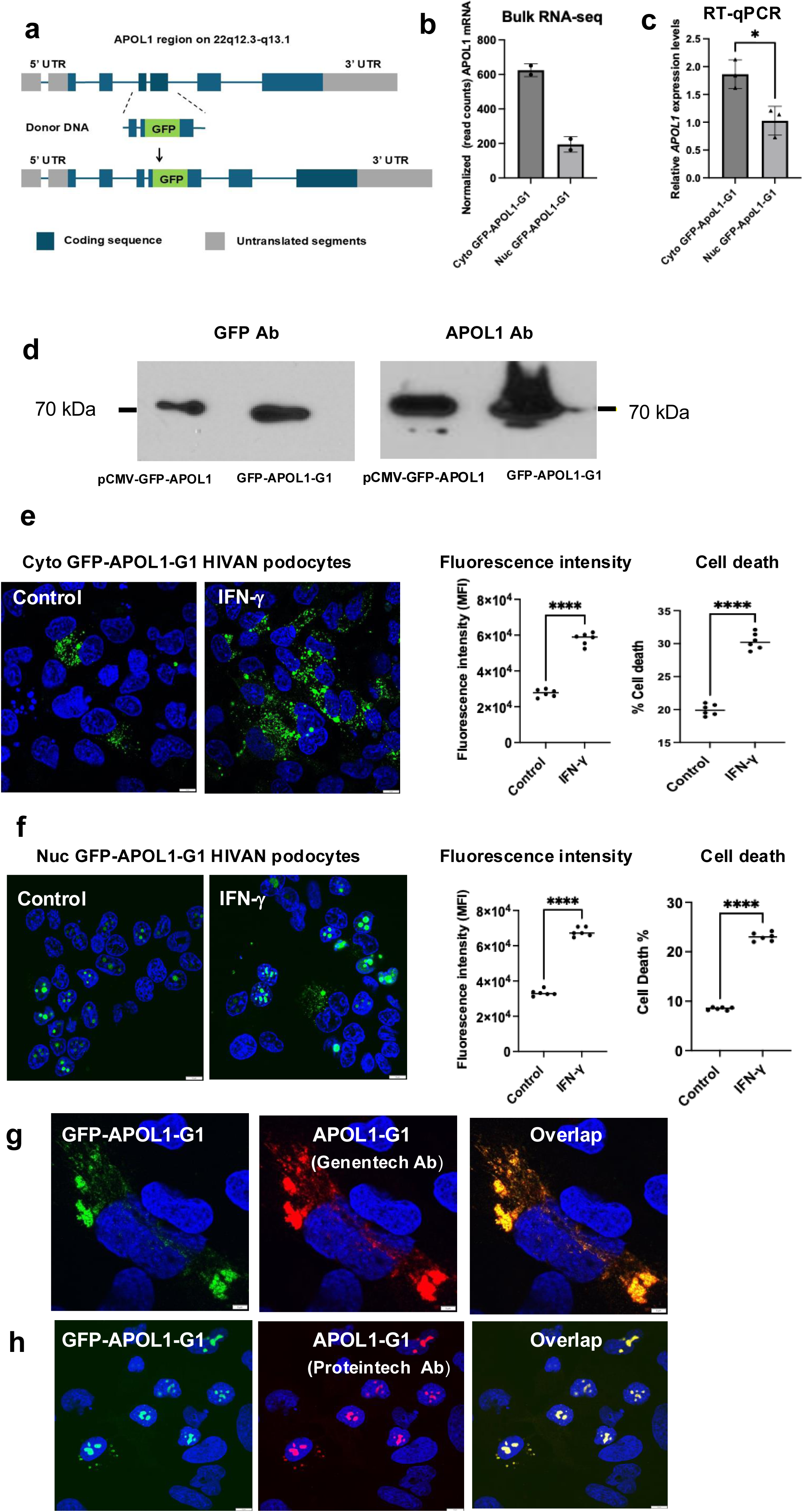
Generation and characterization of cytoplasmic and nuclear APOL1-G1 HIVAN podocyte models. **(a)** Schematic representation of the GFP knock in CRISPR/Cas9-mediated insertion of a GFP gene into the endogenous *APOL1* locus at chromosome 22q13.1. GFP gene was inserted into the APOL1 genomic coding sequence in frame to enable visualization of APOL1 expression by fluorescence microscope. Blue boxes indicate coding regions and gray boxes indicate untranslated regions (UTRs). **(b-c**) Relative APOL1 mRNA expression measured by bulk RNA sequencing and RT-qPCR in HIVAN podocytes expressing cytoplasmic GFP-APOL1-G1 (Cyto GFP–APOL1-G1) or nuclear GFP-APOL1-G1 (Nuc GFP-APOL1-G1). By RT-qPCR, APOL1 expression was significantly higher in Cyto GFP–APOL1-G1 cells. Similar results were obtained by unbiased bulk RNA sequencing of HIVAN podocytes isolated by flow cytometric sorting based on GFP expression. Data are mean ± SD (bulk RNA seq: n= 2 independent biological replicates per group); RT-qPCR; n = 3 independent biological replicates per group). *P* < 0.05. (d) Representative immunoblots of cytoplasmic GFP-APOL1-G1 (Cyto GFP-APOL1-G1) HIVAN podocytes and parental HIVAN APOL1 podocytes transfected with the GFP-APOL1-G1 plasmid. Immunoblotting with antibodies against GFP and APOL1 detected a ∼70-kDa fusion protein corresponding to GFP-tagged APOL1-G1 (GFP, ∼27 kDa + APOL1-G1, ∼42 kDa). Blots were probed with either anti-GFP and anti-APOL1 antibodies from Origene and Sigma-Aldrich respectively, as indicated. Data are representative of *n* = 3 independent experiments. (e) Representative confocal images of cytosolic GFP-APOL1-G1 HIVAN podocytes under basal conditions or following IFN-γ stimulation. Nuclei were counterstained with DAPI. Quantification of fluorescence intensity and cell death of GFP positive cells assessed by flow cytometry is shown in the graphs. Data are mean ± SD from independent experiments (*n* = 6). Scale bars, 10 μM \*\*\*\**P* < 0.0001 using two-tailed Welch’s t test (f) Representative confocal images of nuclear GFP–APOL1-G1 HIVAN podocytes under basal conditions or following IFN-γ stimulation. Nuclei were counterstained with DAPI. Quantification of fluorescence intensity and cell death of GFP positive cells assessed by flow cytometry is shown in the graphs. Data are mean ± SD. from independent experiments (*n* = 6). Scale bars, 10 μM. \*\*\*\**P* < 0.0001 using two-tailed Welch’s t test (g) Representative confocal images of cytoplasmic GFP-APOL1-G1 HIVAN podocytes stained with an anti-APOL1 monoclonal antibody (Genentech), demonstrating co-localization of GFP-tagged APOL1-G1 with APOL1 signal. Nuclei were counterstained with DAPI. Scale bars, 5 μM (h) Representative confocal images of nuclear GFP-APOL1-G1 HIVAN podocytes stained with an anti-APOL1 antibody (Proteintech) raised against the N terminal sequence of APOL1, confirming nuclear accumulation of GFP-APOL1-G1. GFP APOL1-G1 was also detected in the cytosol with the APOL1 antibody. Scale bars, 10, μM

To determine whether GFP insertion altered APOL1 localization, we compared the subcellular distribution of endogenous GFP-APOL1 in knock-in podocytes with the endogenous APOL1 localization in primary HIVAN podocytes. GFP-APOL1 closely recapitulated the intracellular distribution of endogenous APOL1, indicating that insertion of GFP into exon 5 did not appreciably alter APOL1 localization (Fig. 1d–g and Fig. 2e).

Live-cell imaging of APOL1-GFP knock-in podocytes revealed substantial cell-to-cell heterogeneity in APOL1 expression. In the majority of cells, APOL1 exhibited a predominantly endoplasmic reticulum-associated distribution with a prominent perinuclear localization pattern (Fig. 2 e, g), consistent with the localization observed in primary HIVAN podocytes (Fig. 1 e–g). GFP-APOL1-G1 also exhibited a similar association with ER compartments and a comparable distribution between lipid rafts and non-raft membrane fractions (Supplementary Fig. 4). Together, these findings validate the GFP-APOL1-G1 knock-in HIVAN podocyte line as a faithful reporter for monitoring endogenous APOL1-G1 localization.

Unexpectedly, approximately 32% of independently derived APOL1-GFP knock-in clones (8 out of 25) displayed a strikingly different localization pattern characterized by predominant nuclear accumulation of GFP-APOL1-G1 and minimal cytoplasmic staining (Fig. 2f). These nuclear clones expressed lower levels of *APOL1* mRNA than the clones exhibiting the conventional cytoplasmic distribution (Fig. 2 b-c). IFN-γ treatment increased GFP-APOL1 abundance in both cytosolic and nuclear GFP-APOL1-G1 HIVAN podocytes and accelerated podocyte death (Fig. 2 e, f), mimicking the situation of primary podocyte. In cytoplasmic clones, GFP-APOL1-G1 was detected using a specific APOL1 antibody generated by Genentech and raised against the C terminal region of APOL1 (Fig. 2 g) predominately in a perinuclear ER distribution using the ER marker RFP-KDEL (Fig. 3a). In contrast nuclear GFP-APOL1 clones exhibited minimal ER localization and prominent nuclear accumulation of GFP-APOL1 (Fig. 2h) detected with an APOL1 antibody from Proteintech generated against the N terminal region of APOL1. Nuclear clones also showed significant nucleolar staining (Fig. 3b).

**Figure 3.**
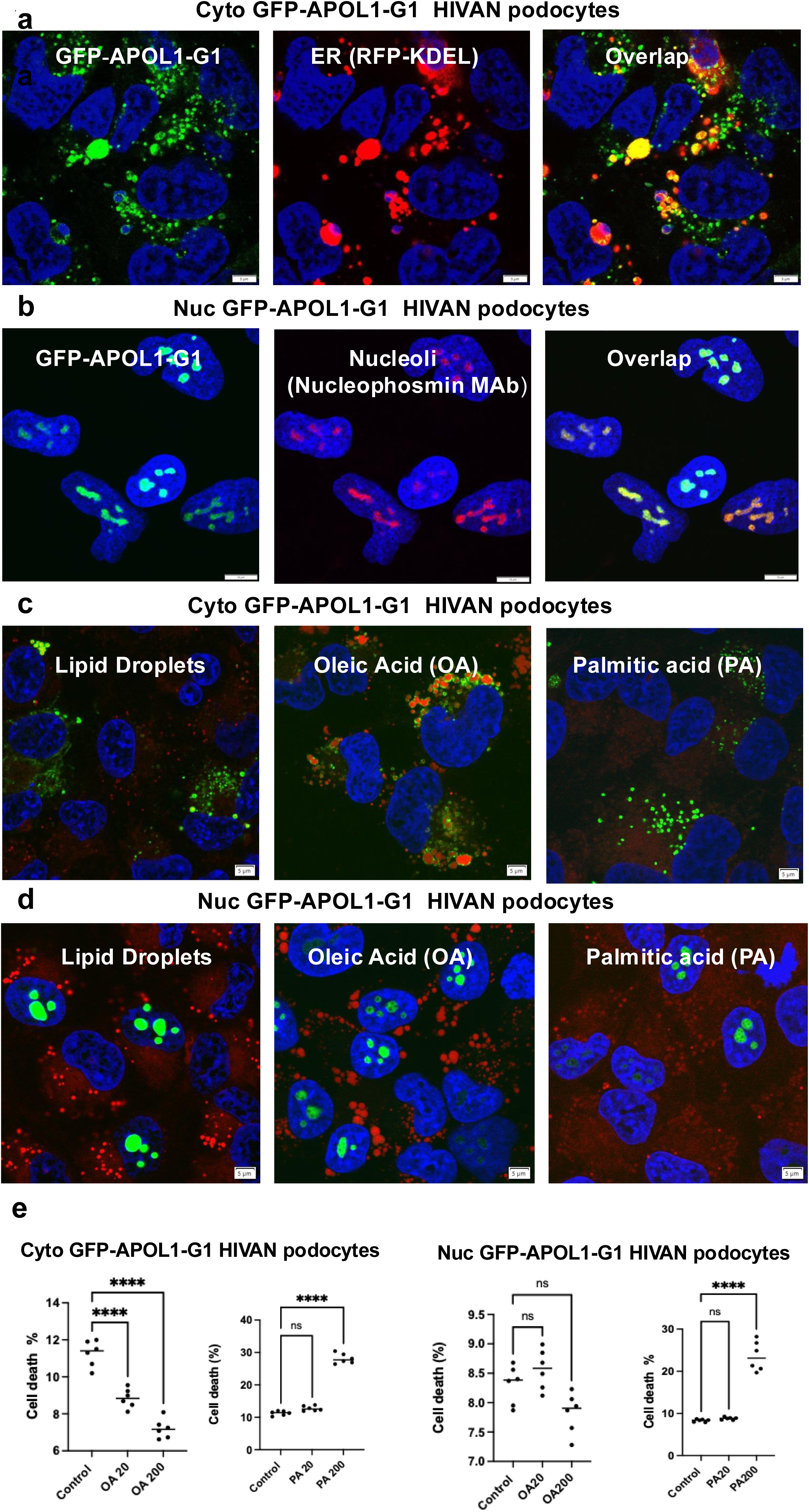
Subcellular localization of APOL1-G1 and modulation of APOL1-associated cytotoxicity by fatty acids in cytosolic and nuclear GFP-APOL1-G1 HIVAN podocytes. (a) Representative confocal images of cytoplasmic GFP-APOL1-G1 HIVAN podocytes co-expressing the endoplasmic reticulum (ER) marker RFP-KDEL. Partial overlap between APOL1-G1 and ER structures is observed. Scale bars, 5 μM. (b) Representative confocal imagens of nuclear GFP-APOL1 G1 HIVAN podocytes co-expressing the nucleoli marker nucleophosmin. Partial overlap between APOL1-G1 and nucleoli structures is observed. Scale bars, 10 μM. (c) Representative confocal images of cytoplasmic GFP-APOL1-G1 HIVAN podocytes under basal conditions or exposed to oleic acid (OA) or palmitic acid (PA). Lipid droplets were stained red with BODIPY 664/676. Cells treated with oleic acid show recruitment of GFP-APOL1 in lipid droplets (overlapping orange color). Results are representative of six independent experiments, Scale bars, 5 μM. (d) Representative confocal images of nuclear GFP-APOL1-G1 HIVAN podocytes under basal conditions or exposed to oleic acid (OA) or palmitic acid (PA). Lipid droplets were stained red with BODIPY 664/676. Oleic acid did not affect the distribution of nuclear GFP-APLO1-G1. Scale bars, 5 μM. (e) Quantification of cell death in cytoplasmic and nuclear GFP-APOL1-G1 HIVAN podocytes following treatment with oleic acid (OA: 20 and 200 μM) or palmitic acid (PA: 20 and 200 μM). OA reduced GFP-APOL1-G1-associated cytotoxicity only in cytoplasmic GFP-APOL1-G1 HIVAN podocytes, whereas high concentrations of PA (200 μM) increased the cytotoxicity in both cytoplasmic and nuclear GFP-APOL1-G1 HIVAN podocytes. Data are presented as mean ± SD from independent experiments (*n* = 6). Each symbol represents an independent biological replicate. ns (non-significant), **** P<0.0001 using one way ANOVA, with Turkey’s multiple comparison test.

### Nuclear APOL1-G1 is resistant to lipid droplet sequestration and promotes podocyte injury

Because APOL1 associates with lipid droplets, we examined whether lipid loading alters APOL1 localization and toxicity. In cytoplasmic GFP-APOL1-G1 podocytes, oleic acid (OA) induced robust recruitment of APOL1-G1 to ADRP/perilipin-2-positive lipid droplets and significantly improved cell survival (Fig. 3c, e), as previously reported by others in non-HIVAN podocytes.^32^ In nuclear GFP-APOL1-G1 HIVAN podocytes, oleic acid did not alter the localization and toxicity of nuclear GFP-APOL1 (Fig. 3 d, e). These findings suggest that the nuclear localization and toxicity of GFP-APOL1-G1 is mechanistically distinct from ER-associated GFP-APOL1-G1 toxicity and is not mitigated by lipid droplet sequestration. In contrast, high concentrations of palmitic acid increased the cell death of both Cyto and Nuc GFP-APOL1-G1 HIVAN podocyte cell lines (Fig 3 c, d, e).

### Nuclear APOL1-G1 activates inflammatory transcriptional programs and directly promotes podocyte death

To define transcriptional pathways associated with nuclear APOL1-G1 expression, we performed RNA sequencing on podocytes expressing predominantly nuclear or cytoplasmic GFP-APOL1-G1. Despite lower total APOL1 transcript abundance (Fig 2b -c), nuclear GFP-APOL1-G1 HIVAN podocytes exhibited a more significant enrichment of IFN-γ response, TNF-α signaling, IL-2/JAK-STAT signaling, inflammatory response, and apoptosis-related pathways, when compared to the Cyto GFP-APOL1-G1 HIVAN podocytes (Fig. 4a–d). Gene set enrichment analysis demonstrated significant activation of the INF-γ, TNF-α IL-6 JAK/STAT and inflammatory pathways among many others (Fig 4 a-d). Gene set enrichment analysis demonstrated significant activation of the INF-γ (NES = 3.01; FDR = 8.3e-10), TNF-α (NES = 2.13; FDR = 1.39e-9), IL-6 JAK/STAT (NES = 1.76; FDR = 0.001), and inflammatory pathways (NES = 2.03; FDR = 2.86e-7) among many others (Fig 4a-d).

**Figure 4.**
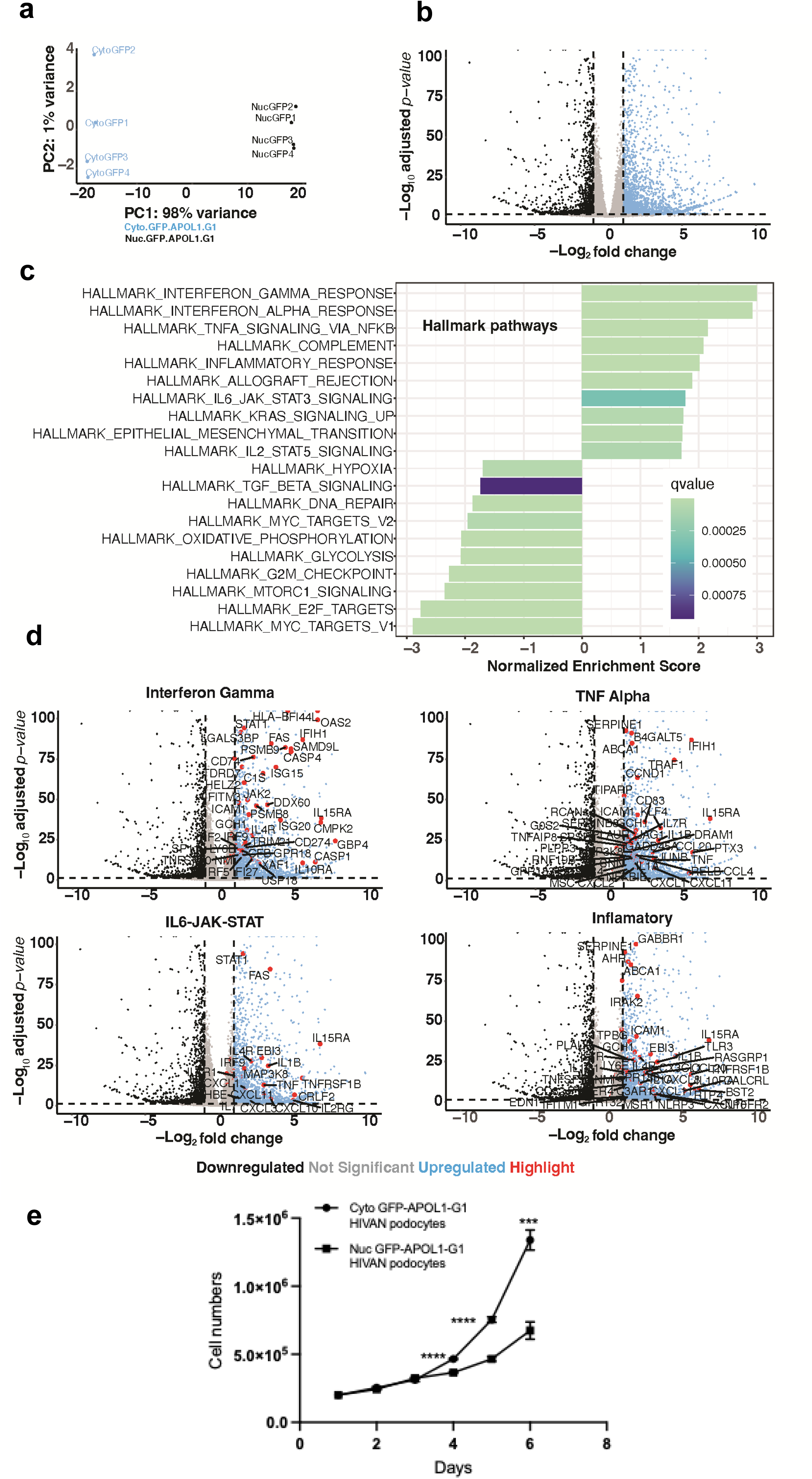
Nuclear localization of APOL1-G1 remodels inflammatory transcriptional programs. **(a)** Principal component analysis (PCA) of RNA-sequencing data from cytoplasmic and nuclear GFP-APOL1-G1 HIVAN podocytes showing clear segregation of transcriptional profiles. **(b)** Volcano plot displaying differentially expressed genes between nuclear and cytoplasmic APOL1-G1 podocytes. Differentially expressed transcripts were defined using [log2fold change +/-1 and adjusted p-value <=0.01]. **(c)** Gene set enrichment analysis of Hallmark pathways. Nuclear GFP-APOL1-G1 expression was associated with enrichment of interferon-γ response, interferon-α response, TNF-α signaling via NF-κB, inflammatory response, IL-6-JAK-STAT3 signaling, and related immune pathways when compared to cytosolic GFP-APOL1-G1 expression. Negative enrichment scores indicate pathways reduced in nuclear APOL1-G1 cells. **(d)** Volcano plots highlighting genes contributing to interferon-γ, TNF-α, IL-6-JAK-STAT, and inflammatory signaling signatures. Representative genes are annotated. RNA-sequencing was performed on independent biological replicates (*n* = 4 per group). Data are presented as mean ± SD where applicable. **(e)** Growth curves of cytoplasmic and nuclear GFP-APOL1-G1 HIVAN podocytes over time. Cell numbers were quantified at the indicated time points (n = 3 per time point). Data are presented as mean ± SD. *** P< 0.001; **** P<0.0001 using two-tailed unpaired Student’s *t*-test.

Nuclear GFP APOL1-G1 podocytes proliferated more slowly than cytoplasmic GFP-APOL1-G1 cells and exhibited more significant spontaneous cell death after seven days in culture (Fig. 4e). To determine whether nuclear localization was sufficient to enhance toxicity, we generated GFP-APOL1-G1 constructs containing a nuclear localization sequence (NLS) and expressed them in APOL1-KO HIVAN podocytes (Fig. 5 a-b and Supplementary Fig 5). Forced nuclear targeting significantly increased cell death compared with non-nuclear targeted APOL1-G1 constructs (Fig. 5c) establishing the nucleus as a direct site of APOL1-mediated injury.

**Figure 5.**
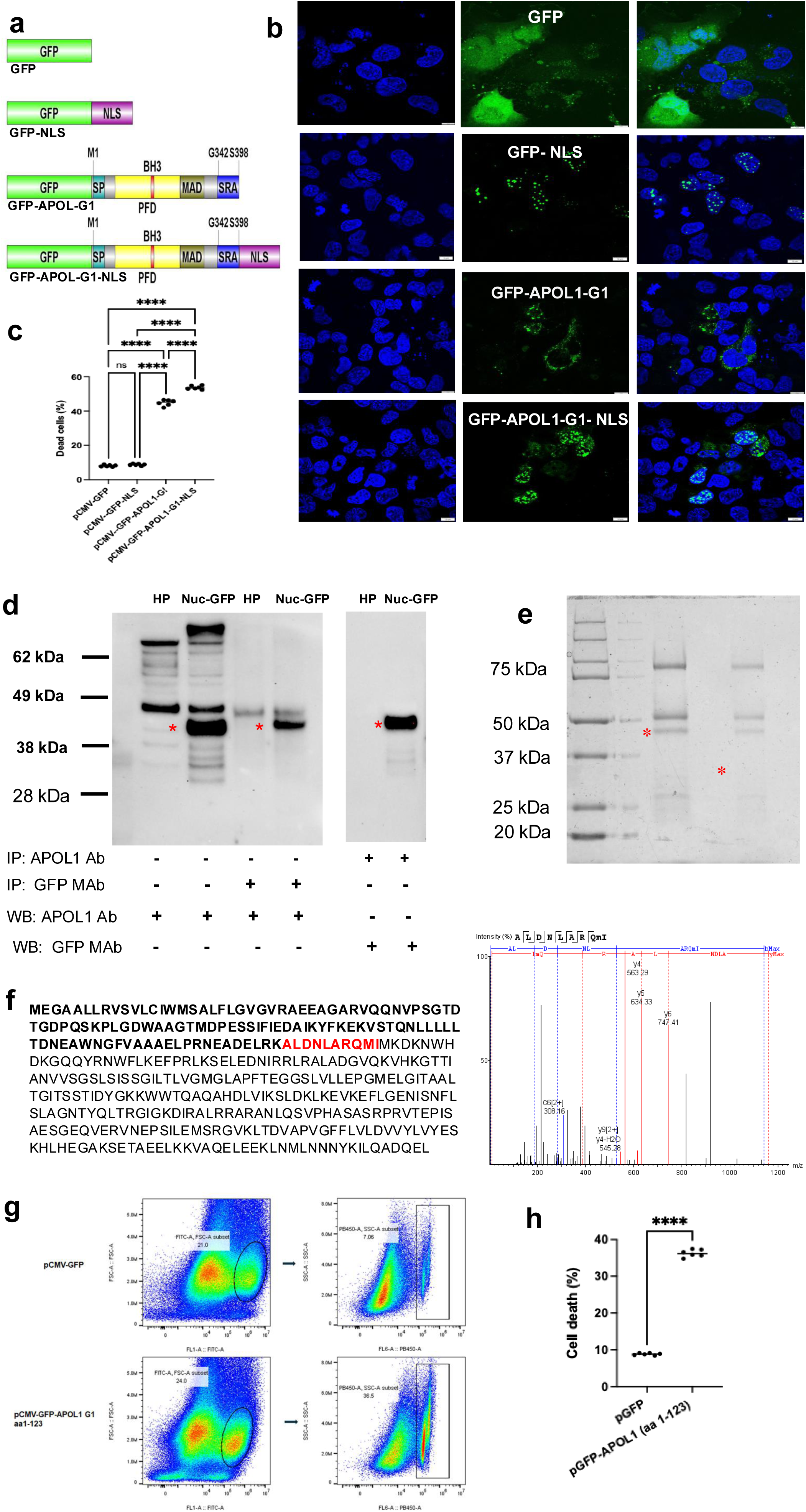
Proteolytic processing of APOL1 generates fragments associated with nuclear toxicity. (a) Schematic representation of GFP-tagged expression constructs, including; GFP control, (CMV-GFP), nuclear localization signal GFP control (CMV-GFP-NLS), GFP-tagged APOL1-G1 (CMV-GFP-APOL1-G1), and nuclear-targeted GFP-APOL1-G1 (GFP-APOL1-G1-NLS). (b) Representative confocal microscopy images showing subcellular localization of the indicated GFP, GFP-NLS, GFP–APOL1-G1, and GFP–APOL1-G1-NLS constructs in APOL1-KO HIVAN podocytes transiently transfected with these constructs. GFP fluorescence (green) and DAPI nuclear staining (blue). Scale bars, 10 μM (c) Forced nuclear localization of GFP-APOL1-G1 significantly increased cytotoxicity compared to GFP-APOL1-G1 lacking an exogenous nuclear localization sequence. The graph shows quantification of cell death in APOL1-KO HIVAN podocytes transfected with the corresponding expression constructs. Data are presented as mean ± SD; ns (non-significant) **** P < 0.001 using one way ANOVA, with Turkey’s multiple comparison test. (d) Co-immunoprecipitation analyses in parental HIVAN podocytes and nuclear GFP-APOL1-G1 cells demonstrating APOL1-G1 N terminal fragments detected using APOL1 antibodies from Proteintech and GFP antibodies from Origene, Asterisks denote APOL1-derived bands subjected to downstream analysis. (e) Representative gel showing excised protein bands analyzed by mass spectrometry. Asterisks indicate fragments selected for peptide identification. (f) Representative tandem mass spectrometry spectrum corresponding to the APOL1-derived nuclear peptide. APOL1 protein sequence highlighting the GFP + APOL1 fragments (aa 1-123). The C-terminal residues of this fragment identified by mass spectrometry are indicated in red. (g) Flow cytometry-based assessment of cell death in APOL1-knockout HIVAN podocytes transfected with pCMV-GFP control or pCMV-GFP–APOL1-G1 aa1–123 plasmids. Twenty-four hours after transfection, cells were harvested, stained with DAPI, and analyzed by flow cytometry. GFP-positive cells were identified by gating on FITC fluorescence and forward scatter parameters, and cell death was quantified as the percentage of DAPI-positive events within the GFP-positive cells. Representative flow cytometry plots are shown. Numbers indicate the percentage of events within each gate. Data were generated from six biological replicates per group (n = 6) in each experiment. The experiment was independently performed twice with similar results. (h) Quantification of cell death in APOL1 KO HIVAN podocytes transfected with either the pCMV-GFP control plasmid or pCMV-GFP-APOL1 aa 1-123 plasmid. Each symbol represents an independent biological replicate. **** P<0.0001 using two-tailed Welch’s t test.

### APOL1 undergoes proteolytic processing to generate distinct N-terminal and C-terminal fragments

To investigate the molecular basis of nuclear APOL1 accumulation, nuclear lysates from GFP-APOL1-G1 podocytes were subjected to immunoprecipitation with APOL1 and GFP antibodies. These studies identified an approximately 42-kDa nuclear fragments composed of GFP fused to the N-terminal 123 amino acids of APOL1 (Fig. 5 d-f). This fragment was detected exclusively by antibodies recognizing the APOL1 N terminus (Proteintech antibody) (Fig. 2h; Supplementary Fig. 6) but not by specific antibodies directed against the C-terminal region from Genentech (data not shown). In addition, the N-terminal fragment of 123 aa alone, induced cell death in APOL1-KO HIVAN podocytes (Fig. 5 g-h). In contrast, IFN-γ-stimulated parental HIVAN podocytes generated an approximately 28-kDa APOL1 fragment detected by the C-terminal-specific APOL1 antibodies from Genentech (Fig. 1 i-j). The appearance of this fragment coincided with the IFN-γ-mediated induction of full-length APOL1 expression and was absent in APOL1-KO HIVAN podocyte exposed to IFN-γ (Fig.1j). Together, these findings demonstrate that APOL1-G1 undergoes proteolytic processing to generate distinct N-terminal and C-terminal cytotoxic fragments that localize to different intracellular compartments.

### Nuclear APOL1-G1 induces a cathepsin-dependent proteolytic program

Differential expression analysis identified significant upregulation of cathepsin S (CTSS), cathepsin L (CTSL), and caspase-3 (CASP3) in nuclear GFP-APOL1-G1 HIVAN podocytes (Fig. 6a) relative to cytosolic GFP-APOL1-G1 HIVAN podocytes. Briefly, CTSS expression increased by 9.55-fold (adjusted P < 1.5e-47), CTSL by 2.2-fold (adjusted P < 6.27e-23), and CASP3 by 1.85-fold (adjusted P < 4.7e-27) relative to cytoplasmic APOL1-G1 HIVAN podocytes (Fig. 6a).

**Figure 6.**
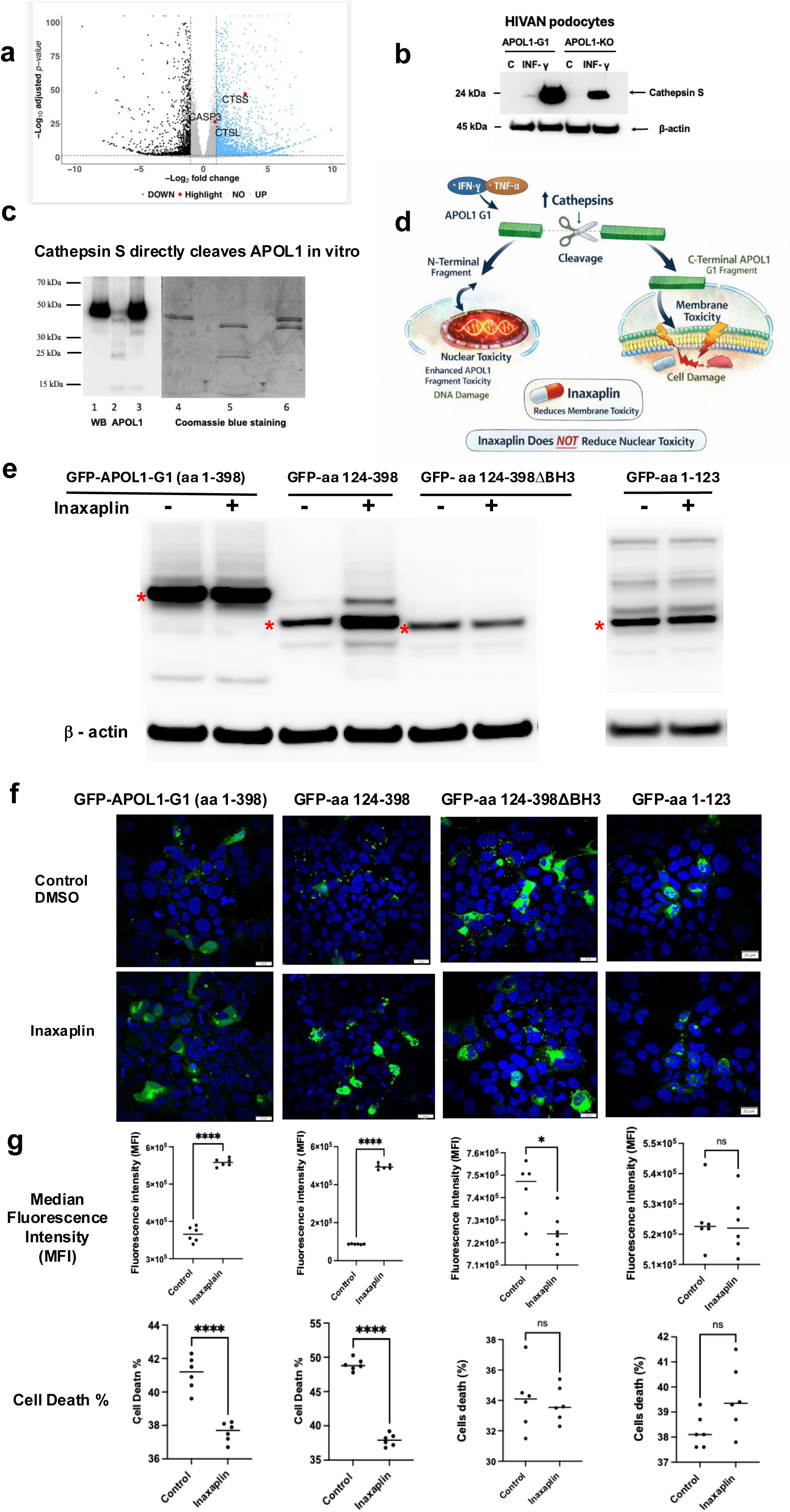
Inflammatory induction of cathepsin S promotes proteolytic processing of APOL1, generating distinct fragments with differential susceptibility to Inaxaplin-mediated protection. (a) Volcano plot indicating the differential gene expression profile between nuclear GFP-APOL1-G1 (Nuc-GFP-APOL1-G1) and cytoplasmic GFP-APOL1-G1 (Cyto-GFP-APOL1-G1) HIVAN podocytes. The x-axis represents log₂ fold change and the y-axis represents −log₁₀ adjusted *P* value. Significantly upregulated genes in nuclear APOL1-G1 podocytes are highlighted in blue, while downregulated genes are shown in black. CTSS (*cathepsin S*) and CTSL (*cathepsin L*) were among the significantly upregulated transcripts in nuclear GFP-APOL1-G1 podocytes, indicating enhanced lysosomal/proteolytic pathway activity associated with nuclear APOL1 localization. CASP3 was also differentially expressed, suggesting activation of cell death–related signaling pathways. Data are representative of 4 independent samples per group. **(b)** Immunoblot analysis of cathepsin S protein expression in HIVAN podocytes, treated with either control vehicle (PBS) or IFN-γ (100 ng/ml) and the corresponding isogenic APOL1-KO HIVAN podocytes, treated in a similar manner. β-actin served as a loading control. The active form of Cathepsin S protein (∼ 24kDa) was detected with the Gene Tex antibody (#GTX638369). Data are representative of two independent biological samples per group. **(c)** Representative APOL1 Western blot analysis of purified recombinant APOL1 (2.5 μg) using the Genentech antibody which recognizes the C-terminal region of APOL1. Lane 1, purified full-length APOL1 control; Lane 2, purified APOL1 incubated with 0.16 μg Cathepsin S; Lane 3, purified APOL1 incubated with 0.16 μg cathepsin S in the presence of 1 mM of the cathepsin S inhibitor. (Apexbio Tech # 1373215-15-6). Cathepsin S treatment resulted in cleavage of APOL1, as evidenced by the appearance of lower-molecular-weight C-terminal fragments and a reduction in the intensity of the full-length APOL1 band. Addition of the cathepsin S inhibitor markedly attenuated APOL1 proteolysis and preserved full-length APOL1. Corresponding Coomassie Brilliant Blue–stained gels (Lanes 4–6) demonstrate the protein species generated under each condition and confirm cathepsin S-dependent cleavage of APOL1. Lane 4, purified full-length APOL1 control; Lane 5 purified APOL1incubated with 0.16 μg cathepsin S; Lane 6, purified APOL1 incubated with 0.16 μg cathepsin S in the presence of 1 mM of the Cathepsin S inhibitor. Data are representative of independent experiments (n=3). **(d)** Proposed model in which inflammatory cytokines induce cathepsin-mediated APOL1-G1 cleavage, generating N-terminal and C-terminal fragments that differentially contribute to nuclear and membrane toxicity. Inaxaplin is proposed to mitigate membrane-associated toxicity but not nuclear toxicity. **(e)** Immunoblot analysis confirming expression of APOL1 truncation and deletion constructs in the presence or absence of Inaxaplin (2.0 μM) in APOL1-KO HIVAN podocytes. β-actin served as a loading control. Red asterisks on the left side indicate the corresponding GFP-APOL1 fusion fragments. Results are representative of two independent experiments. **(f)** Representative confocal images of APOL1-KO HIVAN podocytes expressing the indicated GFP-APOL1 fusion fragments, and treated with control vehicle (DMSO) or Inaxaplin (2.0 μM) Nuclei were stained blue with DAPI. Results were generated with six independent samples in each group. Scale bars, 20 μM. **(g)** Quantification of median fluorescence intensity (MFI) (upper graphs) and cell death (lower graphs) in APOL1-KO HIVAN podocytes transfected with the corresponding APOL1 constructs (shown in Supplementary Fig 7) following exposure to vehicle (DMSO) or Inaxaplin (2.0 μM) Inaxaplin reduced toxicity associated with full-length APOL1-G1 and APOL1-G1 aa124–398 but failed to rescue toxicity induced by constructs lacking the BH3 domain as well as those containing the N-terminal fragment alone. Each pair of cells treated with either DMSO or Inaxaplin was analyzed independently. Data are presented as mean ± SD from independent experiments (*n* = 6). ns (non-significant;)* P<0.05; ****P < 0.0001 using a two tailed Welch’s t test.

Consistent with these transcriptomic findings, IFN-γ markedly increased the expression of the active form of cathepsin S protein in HIVAN podocytes (Fig 6b). Cathepsin S expression was reduced in APOL1-KO HIVAN podocytes and only partially restored by IFN-γ treatment, suggesting functional coupling between APOL1 expression and cathepsin activation (Fig 6b).

To determine whether APOL1 is a direct substrate of cathepsin S, purified recombinant APOL1 was incubated with recombinant cathepsin S. Cathepsin S generated three discrete APOL1 cleavage products (Fig 6c). Immunoblotting using a C-terminal APOL1 antibody demonstrated loss of full-length APOL1 and generation of three C-terminal fragments following cathepsin S treatment (Fig. 6c). Addition of a selective cathepsin S inhibitor largely preserved full-length APOL1 and prevented fragment formation. Coomassie staining independently confirmed cathepsin S-dependent cleavage and its inhibition by pharmacological blockade (Fig. 6c). Together, these findings demonstrate that APOL1 undergoes proteolytic processing to generate distinct N-terminal and C-terminal fragments that localize to different intracellular compartments.

In addition, these observations led us to hypothesize that cathepsin-mediated APOL1 cleavage separates nuclear and membrane toxicities into distinct functional fragments (Fig. 6d). Under this model, inaxaplin (VX-147), a small-molecule inhibitor of APOL1 channel function currently in clinical trials for APOL1-associated kidney diseases,^33,34^ would be expected to inhibit APOL1-mediated membrane toxicity while not affecting the nuclear APOL1 toxicity. We next directly tested this hypothesis.

### Inaxaplin selectively modulates the accumulation and toxicity of APOL1 cleavage products

To determine whether inaxaplin differentially affects full-length APOL1 and APOL1 cleavage products, APOL1-KO HIVAN podocytes were transfected with full-length APOL1-G1, the C-terminal fragment (aa124-398), a mutant C-terminal fragment (aa124-398 ΔBH3) lacking the BH3 domain, or the N-terminal fragment (aa1–123) (Supplementary Fig. 7). We first confirmed the expression levels of each plasmid fragment transfected into APOL1-KO HIVAN podocytes by Western blot analysis (Fig. 6e). We then quantified changes in subcellular localization and fluorescence intensity exclusively between cells transfected with the same fragment, comparing those treated with inaxaplin to those treated with control vehicle (DMSO) (Fig 6 f-g). Inaxaplin produced only a modest increase in full-length APOL1-G1 abundance but markedly increased accumulation of the C-terminal APOL1 fragment (aa124–398) (Fig. 6f-g). The greatest accumulation was observed in cells expressing the C-terminal fragment lacking the BH3 domain (aa124–398 ΔBH3), indicating that the BH3 region restricts intracellular accumulation of APOL1. In contrast, the isolated N-terminal fragment did not exhibit significant inaxaplin-dependent accumulation (Fig. 6 f-g).

Despite increasing intracellular APOL1 abundance, inaxaplin significantly reduced cell death induced by both full-length APOL1-G1 and the C-terminal fragment (aa124–398) (Fig. 6f-g). By contrast, inaxaplin failed to protect podocytes expressing the C-terminal fragment with a BH3 deletion (aa 124-398ΔBH3) as well as the N-terminal fragment (Fig 6 f-g). These findings indicate that intracellular APOL1 accumulation and cytotoxicity can be dissociated and suggest that inaxaplin selectively modulates the accumulation of specific APOL1 fragments.

Overall, these results support a model in which inflammatory signaling induces cathepsin-dependent APOL1 cleavage, generating N-terminal and C-terminal APOL1 fragments with distinct intracellular localizations and biological activities (Fig. 6d). Whereas cathepsin-mediated processing is associated with nuclear APOL1-G1 toxicity, inaxaplin selectively modulates the abundance and cytotoxicity of specific APOL1 fragments, identifying post-translational processing of APOL1 as a potentially targetable pathway in HIV-associated kidney disease.

## Discussion

Our findings identify inflammatory, cathepsin-dependent proteolytic processing of APOL1 as a previously unrecognized mechanism regulating APOL1 biology in human podocytes. Using urine-derived podocytes from children with HIVAN, endogenous CRISPR-engineered APOL1 reporter cell lines, transcriptomic analyses, and biochemical studies, we demonstrate that APOL1 undergoes proteolytic remodeling into distinct protein fragments with divergent intracellular localization patterns and biological activities. APOL1 proteolysis generated nuclear APOL1 fragments associated with inflammatory transcriptional activation, enhanced podocyte injury, and differential responsiveness to the APOL1 inhibitor inaxaplin. Together, these findings suggest that APOL1-associated kidney injury is governed not only by genotype and expression level but also by post-translational processing into biologically distinct protein fragments.

Current models of APOL1 pathogenesis have largely focused on the actions of full-length APOL1. APOL1 localizes to intracellular membranes, including the endoplasmic reticulum, endolysosomal compartments, mitochondria, lipid droplets, and plasma membrane, where it influences ion homeostasis, cellular metabolism, vesicular trafficking, and stress signaling.^6–19^ These observations have motivated therapeutic approaches targeting APOL1 channel activity but do not fully explain the heterogeneity and context dependence of APOL1-associated diseases.^20^ Our findings challenge the prevailing paradigm that full-length APOL1 is the principal pathogenic effector by demonstrating that inflammatory proteolysis generates biologically distinct APOL1 fragments with independent intracellular activities.

The identification of APOL1 proteolysis was enabled by a unique experimental platform combining patient-derived podocytes from children with HIVAN with endogenous CRISPR-engineered APOL1 reporter derivatives. Conventional overexpression systems and available animal models are poorly suited to detect endogenous APOL1 processing, emphasizing the importance of disease-relevant human models for defining APOL1 biology.

The first evidence for APOL1 processing emerged from podocytes derived from the urine of children with HIVAN. These patient-derived podocytes retain expression of developmental markers including CD24 and CD133 while expressing differentiated podocyte markers, recapitulating the dysregulated podocyte state characteristic of HIVAN. Importantly, extensive characterization demonstrated no detectable HIV transcripts or viral proteins, indicating that APOL1 processing occurs independently of ongoing HIV gene expression. Inflammatory stimulation induced robust APOL1 expression together with the appearance of a previously unrecognized ∼28-kDa APOL1 fragment. This fragment was detected in independent HIVAN podocyte lines but was absent in isogenic APOL1-KO cells, arguing against nonspecific degradation or overexpression-related artifacts. These findings suggest that APOL1 processing occurs under physiologic conditions associated with chronic inflammation and kidney injury.

To investigate the significance of this pathway, we generated endogenous GFP-APOL1-G1 reporter HIVAN podocytes. Importantly, N-terminal GFP knock-in did not disrupt the established localization pattern of endogenous APOL1, as most reporter podocytes displayed ER-associated and perinuclear localization consistent with previous reports and studies using validated APOL1 antibodies.^28^ Although most cells exhibited APOL1 localization within these expected compartments, a subset displayed prominent nuclear and nucleolar APOL1 accumulation. These observations suggest that APOL1 trafficking is more dynamic than previously appreciated and that APOL1 may exist in multiple intracellular states with distinct biological consequences.

Nuclear APOL1 accumulation was associated with profound phenotypic changes. Transcriptomic profiling demonstrated enrichment of interferon-response pathways, TNF signaling, IL6-JAK-STAT signaling, inflammatory response programs, and apoptosis-related networks. Nuclear APOL1 podocytes proliferated more slowly and exhibited enhanced spontaneous cell death. Consistent with previous reports^32^, oleic acid promoted recruitment of cytoplasmic APOL1 to lipid droplets and reduced APOL1-mediated cytotoxicity. In contrast, oleic acid neither altered nuclear APOL1 localization nor improved viability in podocytes with predominant nuclear APOL1 accumulation. Forced nuclear targeting experiments further demonstrated that nuclear localization was sufficient to enhance podocyte injury. Together, these findings identify the nucleus as a previously underappreciated compartment of APOL1-mediated injury and support a model in which APOL1 toxicity is partitioned into distinct nuclear and membrane-associated pathogenic programs.

Notably, APOL1 proteolysis and nuclear accumulation were not uniformly observed across all reporter clones. Whereas most clones exhibited predominantly cytoplasmic APOL1 localization, only a subset (32%) developed nuclear APOL1 fragments and associated inflammatory phenotypes. Nuclear APOL1 podocytes also demonstrated increased expression of cathepsins and inflammatory pathways, suggesting that protease activity, inflammatory signaling, lysosomal function, or podocyte differentiation state may influence susceptibility to APOL1 cleavage and nuclear trafficking. These observations raise the possibility that APOL1-mediated injury develops preferentially in specific podocyte states rather than uniformly across all APOL1-expressing cells.

A major conceptual advance arising from this work is the link between APOL1 proteolysis and the emergence of nuclear APOL1 fragments. Proteomic analyses identified an approximately 42-kDa nuclear GFP-APOL1-G1 fragment containing GFP and the N-terminal portion of APOL1, whereas inflammatory stimulation generated inducible ∼28-kDa C-terminal APOL1 fragments in primary HIVAN podocytes. These findings suggest that proteolytic processing expands the biological repertoire of APOL1 by generating fragments capable of accessing distinct intracellular compartments and engaging different pathogenic pathways.

The transcriptional phenotype associated with nuclear APOL1 raises the possibility that APOL1 fragments actively regulate cellular stress responses. Nuclear APOL1 podocytes demonstrated coordinated activation of interferon-responsive genes, inflammatory mediators, and apoptosis-associated pathways. Previous studies have shown that APOL1 can modulate lysosomal stress pathways and TFEB-dependent transcriptional programs,^35^ supporting a broader role for APOL1 in cellular regulation. Whether APOL1 fragments interact directly with chromatin, transcriptional regulators, RNA-processing machinery, or nucleolar structures remains unknown.

Our data further identify cathepsin proteases as candidate mediators of APOL1 processing. Nuclear APOL1 podocytes demonstrated increased expression of CTSS, CTSL, and CASP3, whereas IFN-γ induced robust cathepsin S expression in HIVAN podocytes. Recombinant cathepsin S directly cleaved purified APOL1 in vitro, and pharmacologic inhibition of cathepsin S substantially reduced APOL1 cleavage and preserved full-length APOL1. Together, these findings support a model in which inflammatory signaling activates cathepsin-dependent pathways that remodel APOL1 into distinct protein fragments. Unlike cathepsin L, cathepsin S retains substantial activity at near-neutral pH and can function outside classical lysosomal compartments,^36,37^ raising the possibility that APOL1 processing occurs in multiple intracellular environments.

Although our findings strongly support a role for cathepsins in APOL1 processing, important questions remain. The precise cleavage sites responsible for generating nuclear and cytoplasmic APOL1 fragments have not been fully defined, and it remains unclear whether cathepsin S alone accounts for all cleavage events observed in vivo. Future studies will be needed to map cleavage sites, determine whether APOL1 risk variants alter cleavage susceptibility, and define the molecular targets of nuclear APOL1.

The therapeutic implications of these findings are noteworthy. Inaxaplin was developed on the basis of models centered on full-length APOL1 channel activity.^33,34^ Unexpectedly, we observed that inaxaplin increased intracellular accumulation of specific APOL1 fragments while simultaneously reducing cytotoxicity. Moreover, deletion of the BH3 region resulted in a marked increase in APOL1 protein accumulation, suggesting that this regio contributes to the regulation of APOL1 abundance.

Unexpectedly, inaxaplin failed to significantly attenuate cytotoxicity in HIVAN podocytes expressing the GFP-APOL1 124–398 ΔBH3 construct, despite retaining robust cytoprotective activity against full-length GFP-APOL1-G1 and the GFP-APOL1 124–398 BH3 construct. These findings indicate that increased APOL1 abundance alone is insufficient to predict responsiveness to channel inhibition and suggest that the BH3 region contributes, either directly or indirectly, to the mechanism by which inaxaplin confers cytoprotection. The dissociation between APOL1 accumulation and inaxaplin responsiveness further demonstrates that APOL1 abundance and toxicity can be uncoupled, supporting the concept that inaxaplin primarily modulates the functional state of APOL1 rather than simply reducing its expression.

Several non-mutually exclusive mechanisms may explain these observations. The increased abundance of the GFP-APOL1 aa 124-398 ΔBH3 fragment could reflect altered protein turnover, intracellular trafficking, or conformational stability. Alternatively, deletion of the BH3 region may redirect APOL1-mediated injury toward cytotoxic pathways that are less dependent on the ion-channel activity inhibited by inaxaplin. Because protein stability, channel activity, and drug binding were not directly assessed in the present study, additional mechanistic studies will be required to distinguish among these possibilities. Collectively, these findings extend the established role of the BH3 region in APOL1 toxicity by identifying it as a potential determinant of therapeutic responsiveness to inaxaplin, independent of its effects on APOL1 protein abundance. More broadly, they raise the possibility that therapies targeting membrane-associated APOL1 channel activity may not fully address pathogenic mechanisms mediated by nuclear APOL1 fragments. Together, the differential responses of the GFP-APOL1 aa 124-398 ΔBH3 and GFP-APOL1aa 1-123 fragments indicate that not all APOL1-mediated cytotoxic pathways are susceptible to channel inhibition, highlighting mechanistically distinct modes of APOL1 toxicity.

A major strength of this study is the use of disease-relevant human cellular systems. Because APOL1 is absent from rodents, conventional animal models cannot fully recapitulate endogenous APOL1 biology. ^26^ The concordance of findings across patient-derived podocytes, endogenous reporter systems, transcriptomic analyses, and complementary biochemical studies supports the physiological relevance and robustness of the pathway identified here.

In summary, we identify inflammatory, cathepsin-dependent proteolytic processing of APOL1 as a previously unrecognized mechanism regulating APOL1 biology in disease-relevant human podocytes. APOL1 cleavage generates distinct protein fragments with divergent localization, inflammatory activity, pathogenic potential, and pharmacologic responsiveness. By establishing APOL1 proteolysis as a mechanistically important and therapeutically modifiable process, our findings expand current models of APOL1 pathogenesis and suggest that effective therapies may need to address both membrane-associated and nuclear APOL1 activities.

## Methods

### Human samples

Urine samples were obtained from two groups of African American children living with HIV receiving combination antiretroviral therapy, with HIV-associated chronic kidney disease (HIV-CKD) or preserved kidney function (HIV controls) (n = 8 per group). Groups were matched for age (15 ± 3 years) and sex. Formalin-fixed, paraffin-embedded kidney tissue for immunohistochemical analyses was obtained from renal biopsies or autopsy specimens from individuals with HIV-associated nephropathy (HIVAN) or HIV-positive individuals without kidney disease (n = 4 per group). All studies involving human specimens were approved by the Institutional Review Board of Children’s National Hospital and conducted in accordance with the Declaration of Helsinki.

### Cell culture and generation of isogenic APOL1-edited podocytes

Primary urinary HIVAN podocytes and conditionally immortalized podocyte cell lines were established and characterized as previously described^23,38,39^ and are further described in Supplementary Fig. 1. Cells were maintained in Dulbecco’s modified Eagle medium supplemented with 10% fetal bovine serum, 1 mM L-glutamine, and penicillin–streptomycin at 37 °C in 5% CO₂. Cell identity was confirmed by expression of canonical podocyte markers, and all cell lines were routinely screened by PCR to exclude HIV sequences. Isogenic APOL1 knock-in and knockout cell lines were generated from a previously characterized G1/G1 HIVAN podocyte line ^39^ using CRISPR–Cas9-mediated genome editing. Following puromycin selection, single-cell clones were isolated by limiting dilution and expanded. Correct genome editing and clonal integrity were confirmed by genomic PCR, Sanger sequencing, and whole-genome sequencing. Generation of APOL1-edited podocytes and plasmid construction are described in the Supplementary Methods.

### Immunohistochemistry

Formalin-fixed, paraffin-embedded human kidney sections were subjected to heat-induced antigen retrieval and stained with a monoclonal antibody against APOL1. Immunoreactivity was detected using a biotin–streptavidin–HRP system with tyramide signal amplification and DAB visualization. Detailed staining procedures and antibody information are provided in the Supplementary Methods.

### Immunofluorescence microscopy

Cells grown on poly-L-lysine-coated coverslips were fixed, permeabilized and stained with the indicated primary and fluorescent secondary antibodies. Nuclei and lipid droplets were visualized using DAPI and BODIPY 665/676, respectively. Detailed staining and imaging procedures are provided in the Supplementary Methods.

### Cell-proliferation assay

Podocyte proliferation was assessed by daily cell counting over six days using flow cytometry and a defined number of fluorescent reference cells. Absolute cell numbers were calculated from the ratio of podocytes to reference cells. Additional details are provided in the Supplementary Methods.

### Lipid-raft membrane fractionation

Detergent-resistant membrane fractions were isolated from IFN-γ-treated HIVAN podocytes by discontinuous sucrose-density-gradient ultracentrifugation. Twelve fractions were collected and analyzed by immunoblotting for APOL1 and membrane-compartment markers. Detailed fractionation and immunoblotting procedures are provided in the Supplementary Methods.

#### Western blotting

Cells were lysed in RIPA buffer containing 1% NP-40. Equal amounts of protein were separated by SDS–PAGE and transferred to PVDF membranes. Membranes were blocked, incubated with primary antibodies overnight at 4 °C, followed by HRP-conjugated secondary antibodies, and developed using enhanced chemiluminescence. Signals were detected using Kodak X-OMAT film or a ChemiDoc MP Imaging System (Bio-Rad). Detailed protocols, antibody information, and imaging conditions are provided in the Supplementary Methods.

### Urinary APOL1 quantification

Urinary APOL1 concentrations were measured by enzyme-linked immunosorbent assay (ELISA) and normalized to urinary creatinine (UCr). Detailed assay procedures are provided in the Supplementary Methods.

### RT-qPCR

Total RNA was isolated from cultured cells, reverse transcribed into cDNA, and analyzed by quantitative PCR. Relative APOL1 expression was normalized to GAPDH and calculated using the 2−ΔΔCt method. Primer sequences and PCR conditions are provided in the Supplementary Methods.

### RNA sequencing

Total RNA was isolated from the parental HIVAN podocyte line (P1)^39^ and the corresponding isogenic Cyto GFP-APOL1, Nuc GFP-APOL1 and APOL1-knockout cell (KO) lines. RNA integrity was confirmed before library preparation, and only samples with an RNA integrity number ≥8.0 were sequenced. RNA quality assessment, ribosomal RNA depletion, library preparation and paired-end sequencing (2 × 150 bp) were performed by GENEWIZ (South Plainfield, NJ, USA). Sequencing reads were subjected to quality control, aligned using STAR, and analyzed for differential gene expression using DESeq2. Gene Set Enrichment Analysis (GSEA) was performed to identify enriched biological pathways. Detailed sequencing and computational workflows are provided in the Supplementary Methods.

### Whole-genome sequencing

Whole-genome sequencing was used to confirm CRISPR editing outcomes and assess potential off-target genomic integration of the GFP donor sequence. Briefly, genomic DNA was isolated from cultured podocytes, and whole-genome sequencing libraries were prepared by Novogene (Sacramento, CA, USA) and sequenced on an Illumina NovaSeq platform (2 × 150-bp paired-end reads), generating a mean genome coverage of 31–57×. Sequencing reads were aligned to the hs37d5 decoy human reference genome and analyzed according to GATK Best Practices for germline variant discovery. Variants were annotated using Ensembl Variant Effect Predictor (VEP) and analyzed using GEMINI. Detailed sequencing, quality-control and computational analyses are described in the Supplementary Methods.

### Protein C-terminal sequencing by LC-MS/MS

Nuclear GFP-APOL1-containing protein complexes were isolated by immunoprecipitation, resolved by SDS–PAGE, and subjected to in-gel digestion followed by LC–MS/MS. Peptides were analyzed using a Vanquish Neo nano-LC system coupled to an Orbitrap Fusion Lumos Tribrid mass spectrometer (Thermo Fisher Scientific), and spectra were searched using Byonic software (Protein Metrics). Manual inspection of the MS/MS spectra confirmed that the recombinant GFP–APOL1 protein terminated at amino acid 361, identifying the C-terminal peptide sequence ALDNLARQMI. Detailed mass spectrometry methods are provided in the Supplementary Methods.

### Flow cytometry

Podocytes were treated with recombinant human IFN-γ (10 ng ml⁻¹), transfected with the indicated expression constructs, and treated with vehicle (DMSO) or 2 μM inaxaplin (VX-147), as indicated. Cell viability was assessed by DAPI exclusion, and GFP median fluorescence intensity (MFI) was quantified by flow cytometry. Data were acquired using a CytoFLEX flow cytometer (Beckman Coulter) and analyzed with FlowJo software. Detailed methods are provided in the Supplementary Methods.

### Cathepsin S cleavage assay

Recombinant human APOL1 was incubated with recombinant human cathepsin S in the presence or absence of a cathepsin S inhibitor. Cleavage products were analyzed by SDS-PAGE followed by Coomassie staining or immunoblotting. Detailed methods are provided in the Supplementary Methods.

### Statistics

Data are presented as mean ± s.d. Statistical analyses were performed using GraphPad Prism (v11). Comparisons between two groups were performed using unpaired two-sided Welch’s *t*-tests. Comparisons among three or more groups were performed using one-way ANOVA followed by Tukey’s multiple-comparison test. All statistical tests were two-sided unless otherwise specified. *P* < 0.05 was considered statistically significant. For RNA-sequencing analyses we used Benjamini–Hochberg adjustment and GSEA false-discovery rates. The number of biological replicates (*n*), statistical tests, and exact *P* values are provided in the corresponding figure legends.

## Acknowledgment

The authors gratefully acknowledge Genentech for providing the APOL1 antibodies under a Material Transfer Agreement with Dr. Patricio E. Ray at the University of Virginia. (Scales SJ et al. *J Am. Soc Nephrol*. 2020; 9: 2044-2064).

## Author Contributions

JL and PER designed the experiments. JL, JY, JD, LX, PK, PER, performed experiments and analyzed the data. JL and PER drafted the manuscript. All authors interpreted the results, critically revised and edited the manuscript and approved the final version.

## Data availability

All data supporting the findings of this study are available within the article and its Supplementary Information files. Additional data can be provided by the corresponding author upon request.

