## Supplementary material for "Inflammatory proteolysis generates pathogenic APOL1 fragments with distinct intracellular toxicities in podocytes derived from children with HIV associated nephropathy": S Tables, Figures and legends

### Supplementary Tables

#### Table S1

| Sample | P1 | P1APOL1KO | GFP-APOL1-Cyto | GFP-APOL1-Nuc |
|---|---|---|---|---|
| cov_10X | 0.96 | 0.96 | 0.96 | 0.96 |
| cov_20X | 0.89 | 0.96 | 0.85 | 0.93 |
| cov_30X | 0.64 | 0.94 | 0.54 | 0.79 |
| cov_40X | 0.34 | 0.87 | 0.24 | 0.52 |
| MEDIAN_COVERAGE | 34 | 57 | 31 | 40 |
| MEAN_COVERAGE | 35.2 | 57.3 | 32.1 | 41.06 |
| **SD_COVERAGE** | **15.3** | **19.9** | **14.6** | **16.61** |
| HET_SNP_SENSITIVITY | 0.971031 | 0.972745 | 0.97 | 0.97 |

**Table S1: WGS metrics for all samples** Whole-genome sequencing (WGS) quality metrics for parental HIVAN podocytes (P2), APOL1 knockout podocytes (P2APOL1KO), cytoplasmic APOL1-GFP knock-in clones (GFP-APOL1-Cyto), and nuclear APOL1-GFP knock-in clones (GFP-APOL1-Nuc). Coverage metrics represent the fraction of the genome covered at a minimum depth of 10×, 20×, 30×, or 40× sequencing reads. Median coverage, mean coverage, coverage standard deviation, and heterozygous SNP sensitivity are shown for each sample. These data demonstrate sufficient sequencing depth and coverage for genome-wide assessment of GFP integration sites and structural alterations.

**Table S2**

| sample | total | num_het | num_hom_alt | num_hom_ref |
|---|---|---|---|---|
| P1. APOL1-G1 parenteral podocytes. | 57 | 14 | 43 | 11 |
| GFP-APOL-G1 Cyto HIVAN podocytes | 57 | 12 | 45 | 11 |
| GFP-APOL1-G1 Nuc HIVAN podocytes | 57 | 15 | 42 | 12 |
| APOL1-KO HIVAN podocytes | 65 | 21 | 44 | 4 |

**Table S2: APOL1 Mutation stats**. Summary of APOL1 sequence variants identified by whole-genome sequencing in parental and genome-edited podocyte lines. Total indicates the total number of APOL1 variant sites evaluated. Variant calls were classified as heterozygous (num_het), homozygous alternative (num_hom_alt), or homozygous reference (num_hom_ref). Comparison of APOL1 variant profiles was used to confirm the genetic identity of parental, APOL1 knockout, and APOL1-GFP knock-in podocyte lines and to assess genomic alterations associated with CRISPR-mediated genome editing

### Supplementary Figures.

### Fig. S1.

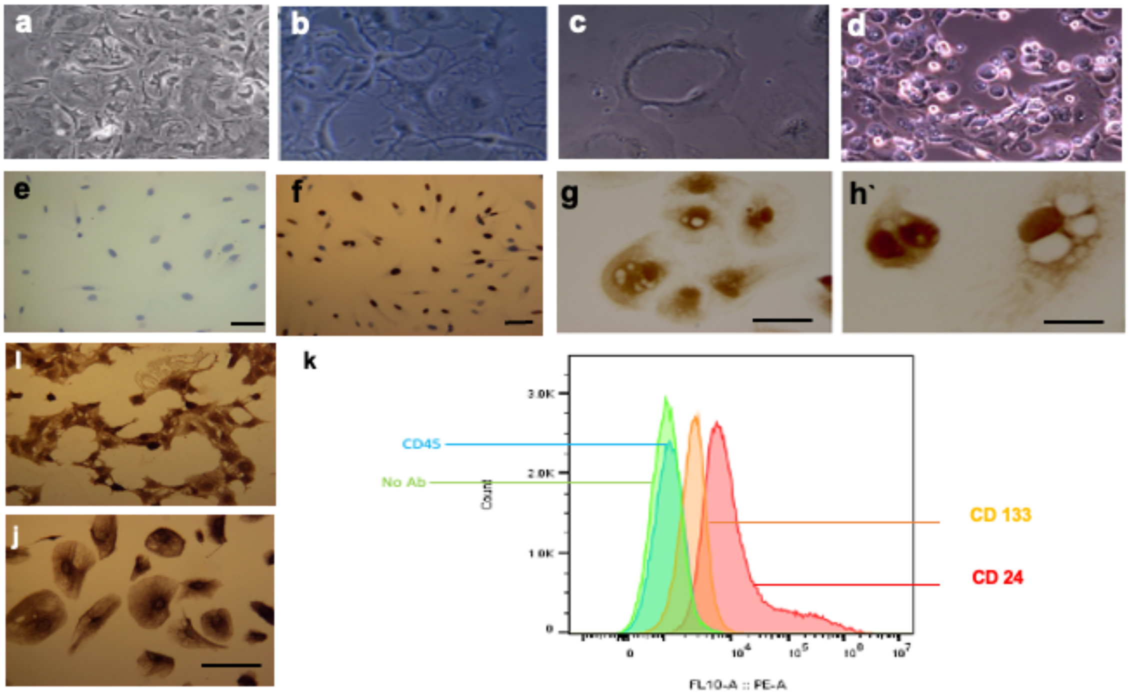


### Fig. S2

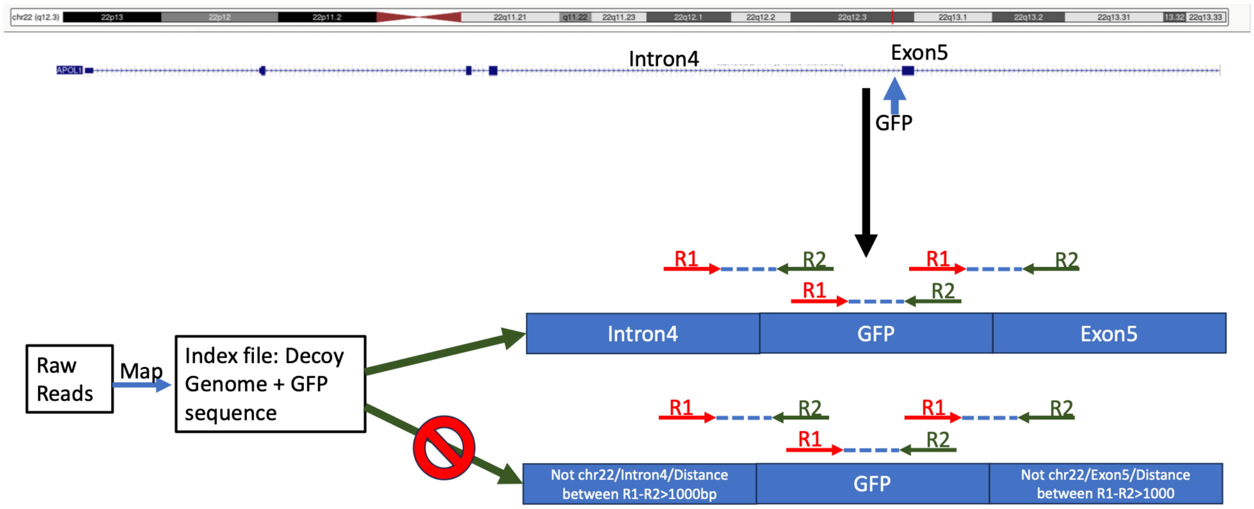

Fig. S3

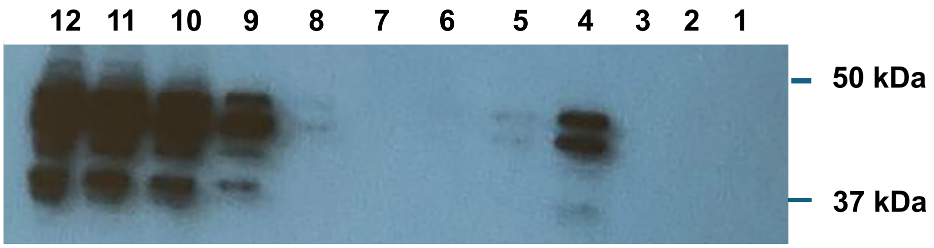
12 11 10 9 8 7 6 5 4 3 2 1
50 kDa
37 kDa

Fig. S4

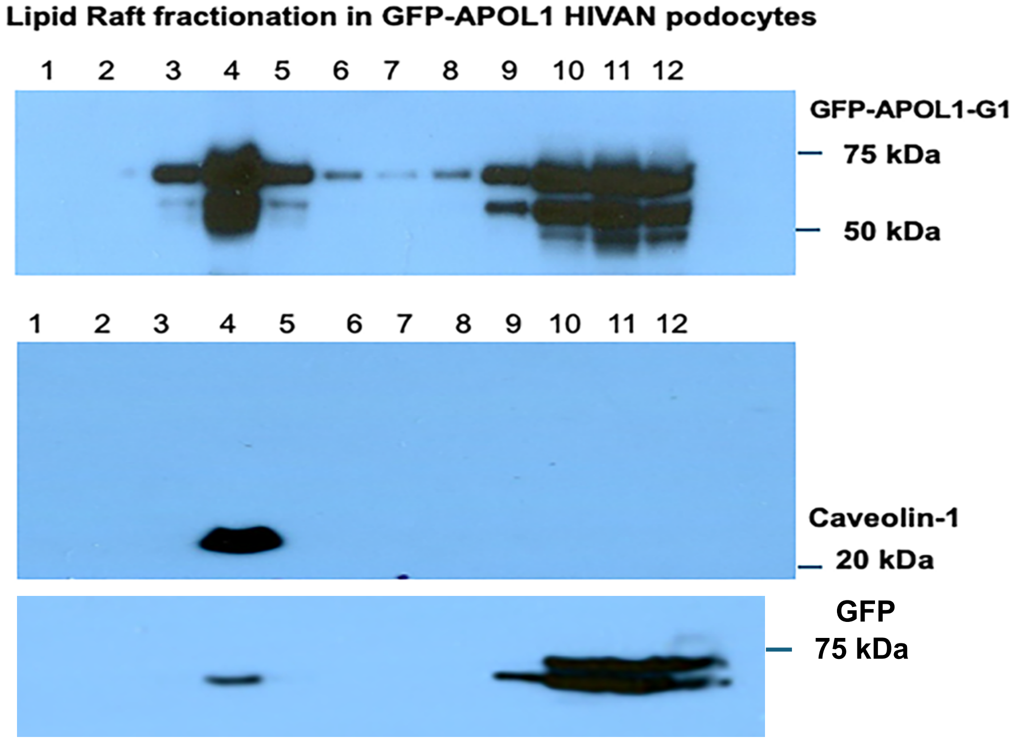
Lipid Raft fractionation in GFP-APOL1 HIVAN podocytes
1 2 3 4 5 6 7 8 9 10 11 12
GFP-APOL1-G1
75 kDa
50 kDa
1 2 3 4 5 6 7 8 9 10 11 12
Caveolin-1
20 kDa
GFP
75 kDa

**Fig. S5**

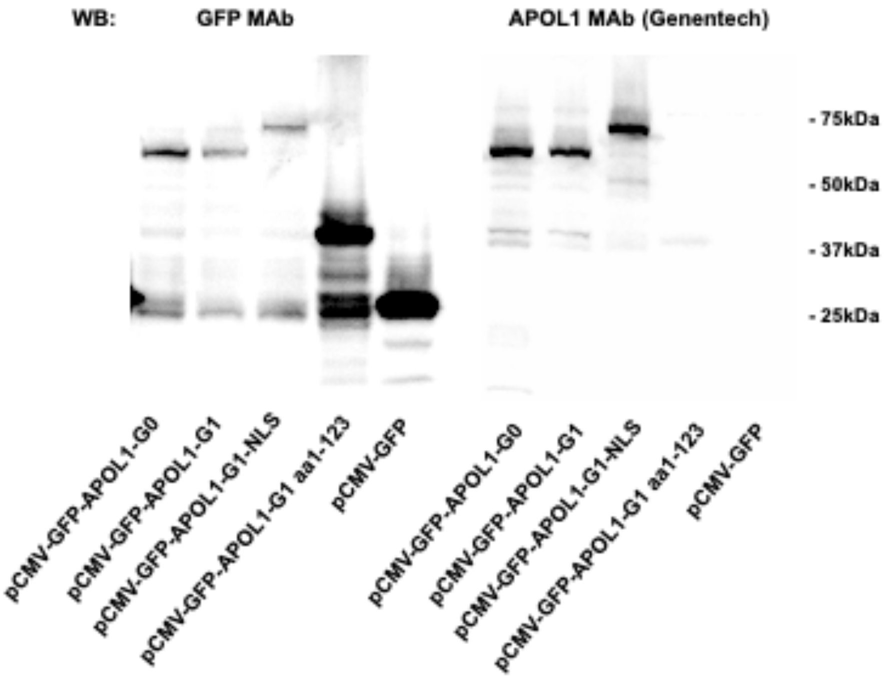
WB:
GFP MAb
APOL1 MAb (Genentech)
- 75kDa
- 50kDa
- 37kDa
- 25kDa
pCMV-GFP-APOL1-G0
pCMV-GFP-APOL1-G1
pCMV-GFP-APOL1-G1-NLS
pCMV-GFP-APOL1-G1 aa1-123
pCMV-GFP
pCMV-GFP-APOL1-G0
pCMV-GFP-APOL1-G1
pCMV-GFP-APOL1-G1-NLS
pCMV-GFP-APOL1-G1 aa1-123
pCMV-GFP

**Fig. S6**

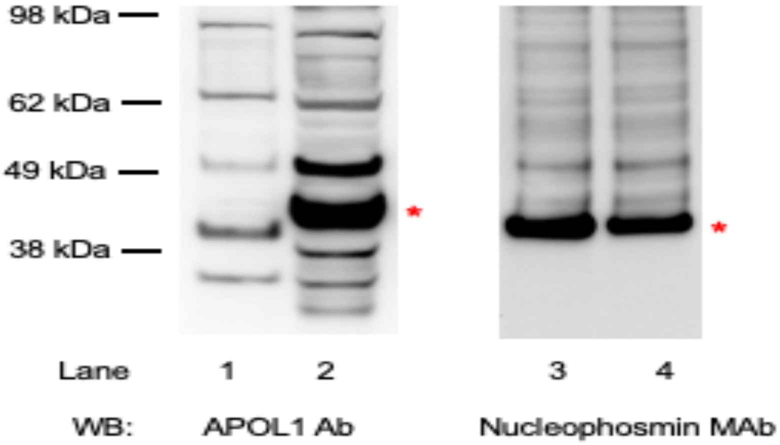
98 kDa
62 kDa
49 kDa
38 kDa
*
*
Lane
1
2
3
4
WB:
APOL1 Ab
Nucleophosmin MAb

**Fig. S7**

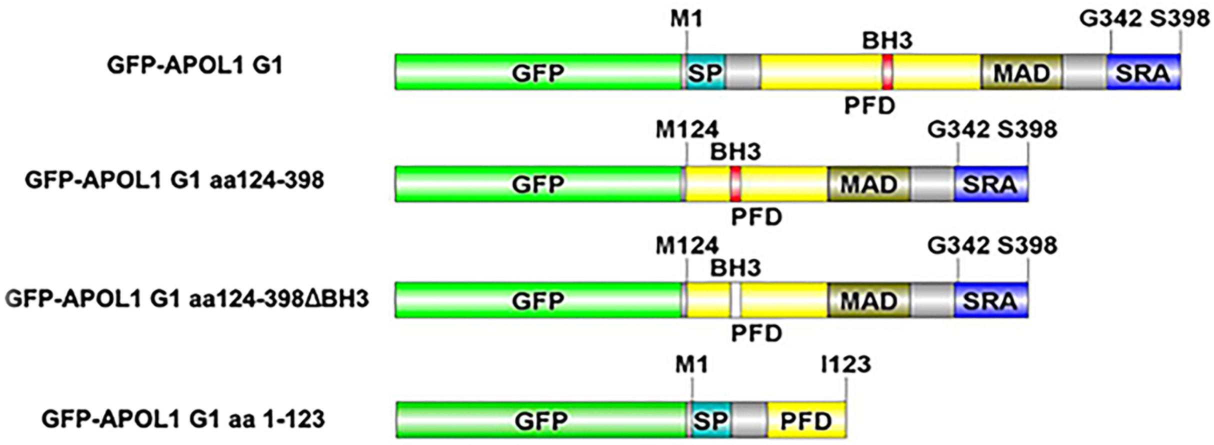
GFP-APOL1 G1
M1
BH3
G342 S398
GFP
SP
MAD
SRA
PFD
GFP-APOL1 G1 aa124-398
M124
BH3
G342 S398
GFP
MAD
SRA
PFD
GFP-APOL1 G1 aa124-398ΔBH3
M124
BH3
G342 S398
GFP
MAD
SRA
PFD
GFP-APOL1 G1 aa 1-123
M1
I123
GFP
SP
PFD

### Supplementary Figure Legends

**Figure S1. Morphological and immunophenotypic characterization of HIVAN podocytes. (a, b)** Representative phase-contrast images of differentiated primary podocytes cultured from the urine of a child with HIV-associated nephropathy (HIVAN). **(c)** Representative image showing vacuolization and senescence of a cultured HIVAN podocyte. **(d)** Representative phase-contrast image of a podocyte cell line transiently transfected with a CMV-driven APOL1-G1 expression plasmid, demonstrating marked cell rounding, detachment, and cell death. **(e, f)** Immunocytochemical staining of HIVAN podocytes incubated with an isotype control antibody **(e)** or an antibody against the podocyte marker WT1 **(f)**, demonstrating nuclear WT1 expression (brown color) in f. **(g)**. Representative dual immunohistochemical staining demonstrating APOL1 expression (brown) and nuclear WT1 expression (red) in HIVAN podocytes. **(i, j)** Representative immunocytochemical staining of HIVAN podocytes for the podocyte-associated marker podocalyxin **(i)** and the epithelial marker KRT18 **(j)**. **(k)** Flow cytometric analysis demonstrating expression of the renal progenitor markers CD24 and CD133 in HIVAN podocytes. No specific staining was detected in the absence of primary antibody (No Ab) or with an antibody against CD45. These findings support a partially differentiated HIVAN podocyte phenotype characterized by the coexistence of podocyte lineage markers and renal progenitor-associated markers. Scale bars, 30 μm.

**Figure S2: Off-Target GFP integration analysis**

Schematic of the strategy used to identify potential off-target integration events in APOL1-GFP knock-in podocytes. Whole-genome sequencing reads were aligned to a custom reference genome containing an additional GFP chromosome. Read pairs containing at least one GFP-mapping read were extracted, and the genomic locations of the corresponding mate reads were analyzed. GFP-associated read pairs mapped exclusively to the APOL1 locus on chromosome 22 at the expected exon 5 knock-in site. No GFP-associated read pairs were detected on other chromosomes or at genomic positions more than 1 kb from the intended insertion site, a distance exceeding the observed library insert size range (150–600 bp). These results provide no evidence for detectable off-target GFP integration events elsewhere in the genome.

**Figure S3. Lipid raft fractionation of HIVAN podocytes transiently transfected with APOL1-G1.**

Membrane fractions were isolated from HIVAN podocytes transiently transfected with APOL1-G1 by detergent-resistant membrane (lipid raft) fractionation. Twelve sequential fractions (1–12) were collected, and equal volumes of each fraction were analyzed by immunoblotting using the Sigma-Aldrich anti-APOL1 rabbit polyclonal antibody (HPA018885). APOL1-G1 was detected predominantly in the higher-density, non-raft fractions (fractions 9–12), with additional localization in the lipid raft fractionS (fractions 5-4).

**Figure S4. Lipid raft fractionation of cytosolic GFP-APOL1-G1 HIVAN podocytes.**

Membrane fractions were isolated from cytosolic GFP-APOL1-G1 HIVAN podocyte cell line by detergent-resistant membrane (lipid raft) fractionation. Twelve sequential fractions (1-12) were collected, and equal volumes of each fraction were analyzed by immunoblotting using the Sigma-Aldrich anti-APOL1 rabbit polyclonal antibody (HPA018885) or the GFP antibody from Origene. GFP-APOL1-G1 was detected predominantly in the higher-density, non-raft fractions (fractions 9–12), with additional localization in the lipid raft fractions (fractions 3–5). Caveolin-1, a lipid raft marker, was enriched in fraction 4, confirming successful isolation of detergent-resistant membrane domains. Molecular weight markers are shown on the right; GFP-APOL1-G1 migrated at approximately 70 kDa, and caveolin-1 migrated at approximately 22 kDa.

**Figure S5 Validation of GFP-APOL1 construct expression in APOL1-knockout HIVAN podocytes by western blot.** APOL1-knockout HIVAN podocytes were transiently transfected with pCMV-GFP-APOL1-G0, pCMV-GFP-APOL1-G1, pCMV-GFP-APOL1-G1-NLS, pCMV-GFP-APOL1-G1 aa1–123, or pCMV-GFP control plasmids. Whole-cell lysates were harvested 24 h after transfection and analyzed by western blot using the Origen anti-GFP antibody (left) or the Genentech anti-APOL1 antibodies (clones 3.1CA80C/3.7D6P80C) used throughout this study (right). GFP-APOL1 fusion proteins and the GFP control were detected at their expected molecular weights. Molecular weight markers are indicated. Shown is a representative blot from two independent experiments with similar results.

**Figure S6 Detection of nuclear GFP-APOL1 in HIVAN podocytes.** Nuclear lysates were prepared from parental HIV-associated nephropathy (HIVAN) podocytes (lanes 1 and 3) and HIVAN podocytes expressing nuclear GFP-APOL1-G1 (Nuc-GFP–APOL1-G1; lanes 2 and 4). Proteins were analyzed by western blot using an antibody directed against the N-terminus of APOL1 (Proteintech; lanes 1–2) and an anti-nucleophosmin antibody as a nuclear marker and loading control (lanes 3-4). A GFP-APOL1-specific band was detected at approximately 42 kDa in nuclear lysates (27 kDa-GFP + ~ 13-15 kDa) N-Terminal fragment of APOL-1 from Nuc-GFP-APOL1-G1-expressing cells (asterisks), but not in parental HIVAN podocytes. Nucleophosmin immunoblotting confirmed enrichment of nuclear proteins in both samples. Shown is a representative blot from two independent experiments with similar results.

**Figure S7. Schematic representation of GFP-tagged APOL1-G1 full-length and deletion constructs.** Schematic illustration of full-length GFP-APOL1-G1 and GFP-APOL1-G1 deletion constructs encompassing amino acid residues 124–398 (aa124–398), aa124–398ΔBH3, and aa1–123. Functional domains are indicated, including the signal peptide (SP), BH3 domain (BH3), pore-forming domain (PFD), membrane-addressing domain (MAD), and SRA-interacting region (SRA). Amino acid boundaries of each construct are shown.

### Supplementary Methods

**Materials and reagents** Synthetic nucleic acid reagents, recombinant proteins, antibodies, plasmids, chemical reagents, and other commercial materials used in this study were obtained from the suppliers listed in **Supplementary Tables 1-3** or in the main text of the manuscript and supplementary information listed below. All restriction enzymes were purchased from New England Biolabs (Ipswich, MA). All reagents were used according to the manufacturers' instructions unless indicated otherwise in the text.

**Supplementary Table 1. Key reagents**

| Reagent | Supplier | Catalog No |
|---|---|---|
| Synthetic nucleic acids | Eurofins Genomics | |
| Oleic Acid | Thermo Fisher Scientific | A195-500 |
| Palmitic Acid | Thermo Fisher Scientific | AC129702-500 |
| DAPI | Thermo Fisher Scientific | D1306 |
| Recombinant human IFN-γ | Thermo Fisher Scientific | PHC4031 |
| Puromycin dihydrochloride | Thermo Fisher Scientific | A1113803 |
| Lipofectamine 3000 | Thermo Fisher Scientific | L3000015 |
| DNAzol | Thermo Fisher Scientific | 10-503-027 |
| Fluoromount-G | Thermo Fisher Scientific | 50-187-88 |
| BODIPY 665/676 | Thermo Fisher Scientific | B3932 |
| Recombinant human APOL1 protein | Sino Biological | 13910-H08B |
| Recombinant human Cathepsin S protein | Sino Biological | 10487-H08H |
| Cathepsin S inhibitor | Apexbio Technology | 1373215-15-6 |
| GoTaq® DNA Polymerase | Promega | M3001 |
| pGL3-Basic Vector | Promega | E1751 |
| pDsRed-Monomer-C1 | Takara Bio | 632466 |
| pZsGreen1-C1 | pZsGreen1-C1 | 632447 |

**Supplementary Table 2. Plasmids**

| Plasmid | Source | Catalog No |
|---|---|---|
| pLentiCRISPRv2 | Addgene (gift from the donor) | 52961 |
| pBFP-KDEL | Addgene (gift from the donor | 49150 |
| pEGFP-ADRP | Addgene (gift from the donor) | 87161 |
| pGL3-Basic | Promega | E1751 |
| pDsRed-Monomer-C1 | Takara Bio | 632466 |
| pZsGreen1-C1 | Takara Bio | 632447 |

**Supplementary Table 3. Antibodies**

| Antibody | Supplier | Catalog No or name | Application |
|---|---|---|---|
| Anti-APOL1 | Millipore Sigma | HPA0 18885 | ICH/WB |
| Anti-APOL1 (N terminal epitope) | Proteintech | 11486-2-AP | IF/IP/WB |
| Anti-APOL1 (gift from donor) ref # 8 | Genentech | name: 5.17d12 | IHC |
| Anti-APOL1 (gift from donor) ref # 8 | Genentech | name: 3.1C1 A80C | WB |
| Anti-APOL1 (gift from donor) ref # 8 | Genentech | name: 3.7D6 P80C | WB |
| PE anti-human CD24 | Bio Legend | 311105 | Flow Cyto |
| PE anti-human CD133 | Bio Legend | 372803 | Flow Cyto |
| PE anti human CD45 | Bio Legend | 368509 | Flow Cyto |
| Goat anti mouse IgG (H+L) Rhodamine | Thermo Fisher Scientific | R-6394 | WB |
| Goat anti Rabbit IgG (H+L) Cross Adsorbed, Rhodamine Red™-X | Thermo Fisher Scientific | 31660 | WB |
| Zs Green1 monoclonal clone OTI2C2 | Origene | TA180002 | IP-WB |
| B23 Nucleophosmin (NA24) | Santa Cruz | Sc53175 | IF |
| Anti-cathepsin (clone HL2302) | Gene Tex | GTX638369 | WB |
| Anti-caveolin 1 | BD Transduction | 640406 | WB |
| Anti-calnexin | Abcan | Ab10286 | WB |
| Mouse monoclonal anti-WT1 (clone 6F-H2) | Agilent/Dako | M3561 | ICH |
| Mouse monoclonal anti-podocalyxin (clone 3D3) | Invitrogen | 39-3800 | ICH |
| Mouse monoclonal nti-KRT18 (CK18) clone (clone DC10) | Abcam | Ab7797 | ICH |
| Anti-β actin | Sigma Aldrich | A1978 | WB |
| HRP-anti-rabbit IgG | Cell Signaling | 7074 | WB |
| Biotinylated donkey anti rabbit IgG (H+L) | Jackson Immune Research | 711-066-152 | WB |
| HRP-Donkey anti mouse IgG (H + L) | Jackson Immune Research | 715-035-151 | WB |

### Human Samples

Urine samples were obtained from two groups of African American children living with HIV receiving combination antiretroviral therapy, with HIV-associated chronic kidney disease (HIV-CKD) or preserved kidney function (HIV controls) (n = 8 per group). The groups were similar in age (approximately 15 ± 3 years) and sex. Urine samples were collected and processed following standard procedures as we described before.[1] Formalin-fixed, paraffin-embedded kidney tissue for immunohistochemical analyses was obtained from renal biopsies or autopsy specimens from individuals with HIV-associated nephropathy (HIVAN) or HIV-positive individuals without kidney disease (n = 4 per group). HIV-CKD, HIVAN, and HIV controls cases were defined using standard definitions as described before.[2] All studies involving human specimens were approved by the Institutional Review Board of Children's National Hospital and conducted in accordance with the Declaration of Helsinki.

### Cell culture and generation of isogenic APOL1-edited podocytes

Primary urinary HIVAN podocytes and conditionally immortalized podocyte cell lines were established and characterized as previously described[3-5] and are further described in Supplementary Fig. 1. Cells were maintained in Dulbecco's modified Eagle medium supplemented with 10% fetal bovine serum, 1 mM L-glutamine, and penicillin–streptomycin at 37 °C in 5% $CO_2$. Cell identity was confirmed by expression of canonical podocyte markers, and all cell lines were routinely screened by PCR to exclude HIV sequences. Isogenic APOL1 knock-in and knockout cell lines were generated from a previously well characterized G1/G1 HIVAN podocyte line described in detail before[3], using CRISPR–Cas9-mediated genome editing.as described before. [6,7] Following puromycin selection, single-cell clones were isolated by limiting dilution and expanded. Correct genome editing and clonal integrity were confirmed by genomic PCR, Sanger sequencing, and whole-genome sequencing as described below.

### Immunohistochemistry

Immunohistochemistry was performed on formalin-fixed, paraffin-embedded kidney sections from individuals with HIV-associated nephropathy and HIV-positive individuals without kidney disease. All four kidney sections per group were deparaffinized in xylene and rehydrated through graded ethanol solutions. Heat-induced antigen retrieval was performed in DAKO Target Retrieval Solution (GV805) at 99 °C for 20 min.

Endogenous avidin and biotin activity was blocked using the Avidin/Biotin Blocking Kit (SP-2001) from Vector Laboratories (Newark CA). Sections were incubated with rabbit monoclonal anti-APOL1 antibody (clone 5-17D12) at 0.5 µg ml⁻¹ for 60 min at room temperature[8], followed by biotinylated donkey anti-rabbit IgG (711-066-152, Jackson ImmunoResearch) at 5 µg ml⁻¹ for 30 min. Signal was detected using streptavidin-HRP for 30 min, followed by tyramide signal amplification (PerkinElmer) for 2–5 min. Immunostaining was visualized with 3,3′-diaminobenzidine (DAB) and sections were counterstained according to standard protocols. WT1, podocalyxin and CK18 were detected using EnVisio HRP polymer system (Dako, K4001) and visualized with DAB. Sections were counterstained with hematoxylin. Human kidney tissue served as the positive control, while omission of the primary antibody and a mouse IgG1 isotype control served as negative controls.

**Immunofluorescence microscopy**

Cells grown on poly-L-lysine-coated coverslips were fixed with 4% paraformaldehyde and permeabilized with 0.1% Triton X-100. Nuclei were stained with DAPI (1 µg ml−1) and lipid droplets were stained using BODIPY 665/676 (1 µg ml-1). For immunostaining, cells were blocked with 2% fetal bovine serum for 1 h and incubated with primary antibodies (0.5 µg ml−1) for 1 h at room temperature. Following washing with PBS containing 0.1% Tween-20, cells were incubated with rhodamine red-conjugated secondary antibodies (1 µg ml−1) and mounted using Fluoromount-G. Imaging was performed using Olympus FV1000 or FV4000 inverted confocal microscopes. Images were acquired with a 60× or 100×/1.40 NA oil-immersion objective using FLUOVIEW Smart operating software. Acquisition settings were kept constant across all samples and the corresponding negative controls. Image analysis was carried out using Olympus FLUOVIEW and OlyVIA software

**Western blotting.** Cells were lysed in RIPA buffer containing 1% NP-40. Equal amounts of protein were resolved on 10%, 12%, or 4–20% Tris-glycine SDS–PAGE gels, selected according to the molecular weight of the target protein. Precision Plus Protein Dual Color Standards (1610374, Bio-Rad) were used as molecular weight markers. Proteins were transferred to PVDF membranes and blocked for 30 min at room temperature in PBS containing 5% (w/v) nonfat dry milk and 0.1% Tween-20. Membranes were incubated overnight at 4 °C with primary antibodies diluted in blocking buffer, washed four times for 10 min with TBST, and incubated for 1 h at room temperature with HRP-conjugated donkey anti-rabbit IgG (711-036-152, Jackson ImmunoResearch) or the appropriate HRP-conjugated secondary antibody. Following four additional washes with TBST, immunoreactive bands were detected using SuperSignal West Pico (34577) or SuperSignal West Femto (34096) chemiluminescent substrate (Thermo Fisher Scientific) according to the manufacturer's instructions. Chemiluminescent signals were captured using Kodak X-OMAT film (Kodak Scientific Imaging) or a ChemiDoc MP Imaging System (Bio-Rad). For detection of APOL1 fragments in APOL1-knockout HIVAN podocytes, 1.5 × $10^6$ cells were seeded in 10-cm dishes and transfected with 10 µg plasmid DNA using 10 µl Lipofectamine. Optimal APOL1 detection was achieved using a combination of the APOL1 monoclonal antibodies 3.1C1 A80C and 3.7D6 P80C (both from Genentech), each at a final concentration of 0.05 µg $ml^{-1}$, anti-cathepsin S (clone HL2302) and anti-β actin.

**APOL1 ELISA.**

Urinary APOL1 concentrations were quantified in urine samples from HIV controls (HIV-C) and individuals with HIV-associated chronic kidney disease (HIV-CKD) using the Human APOL1 ELISA Kit (Proteintech, cat. no. KE00047) (sensitivity 0.07ng/ml; range between 0.156-10 ng/ml) according to the manufacturer's instructions Urinary creatinine (UCr) was measured using the Creatinine Parameter Assay Kit (R&D Systems, cat. no. KGE005), and urinary APOL1 levels were normalized to urinary creatinine (UCr.)

Urine samples were centrifuged at 2,500 rpm for 10 min, and the supernatants were collected for analysis.[1] Samples were diluted 1:10 in distilled water and assayed in duplicate. Absorbance was measured at 450 nm with wavelength correction at 630 nm using a microplate reader. The APOL1 ELISA had a detection sensitivity of 0.07 ng $ml^{-1}$ and a dynamic range of 0.156–10 ng $ml^{-1}$. Urinary creatinine was quantified using the Jaffe reaction according to the manufacturer's protocol. Absorbance was measured at 490 nm, and creatinine concentrations were calculated from a standard curve.

**Lipid raft membrane fractionation**

Lipid raft fractions were isolated from HIVAN podocytes treated with IFN-γ (100 ng $ml^{-1}$) for 24 h. Approximately 5 × $10^7$ cells were lysed on ice in MBS-T buffer (25 mM MES, pH 6.5, 150 mM NaCl, 1% Triton X-100, and protease inhibitor cocktail (Roche; 1 tablet per 50 ml)). Cell lysates were homogenized using a Dounce homogenizer (three strokes), followed by three 20-s bursts with a Polytron homogenizer and a 30-s burst with a probe sonicator (Sonic 300 Dismembrator).

Lysates were mixed with an equal volume of 80% sucrose prepared in MBS buffer, overlaid with 35% and 5% sucrose solutions to generate a discontinuous sucrose gradient (80%:35%:5%, 1:2:1, v/v/v), and centrifuged in an SW55Ti rotor (Beckman Coulter) at 187,813 × *g* (39,000 rpm) for 16 h at 4 °C. Twelve fractions (~300 μl each) were collected sequentially from the top of the gradient. Aliquots (25 μl) of each fraction were analyzed by SDS–PAGE and immunoblotting.

Primary antibodies included anti-APOL1 (clones 3.1CA80C/3.7D6P80C), anti-calnexin (Abcam, ab10286), anti-caveolin-1 (BD Transduction Laboratories, 610406), anti-cathepsin S (GeneTex, GTX638369), and anti-β-actin (Sigma-Aldrich, A1978). Primary antibodies were used at a 1:1,000 dilution. HRP-conjugated donkey anti-mouse IgG (H+L) (Jackson ImmunoResearch) and HRP-conjugated anti-rabbit IgG (Cell Signaling Technology, 7074) were used at a 1:2,000 dilution.

**RT-qPCR.** Total RNA was extracted from cultured cells using the RNeasy Plus Micro Kit (Qiagen) according to the manufacturer's instructions. First-strand cDNA was synthesized from 1 μg of total RNA in a 20-μL reaction using qScript cDNA SuperMix (VWR). Quantitative real-time PCR was performed in technical triplicates using 1 μL of cDNA per reaction with GoTaq qPCR Master Mix (Promega) on a CFX Connect Real-Time PCR System (Bio-Rad). Relative *APOL1* mRNA expression was normalized to *GAPDH* and calculated using the 2−ΔΔCt method. Amplification specificity was confirmed by melt-curve analysis. PCR cycling conditions were 95 °C for 3 min, followed by 40 cycles of 95 °C for 15 s and 60 °C for 1 min. Primer sequences for human *APOL1* were forward 5′-ATAATGAGGCCTGGAACGGAT-3′ and reverse 5′-GGATGCCAGAGGAAATGCTG-3′. *GAPDH* primer sequences have been reported previously.[3]

**CRISPR guide RNA design.**

A guide RNA targeting exon 5 of APOL1 (5′-GTGCAACAAAACGTTCCAAG-3′) was designed using theTrueDesign Genome Editor platform (Thermo Fisher Scientific). Complementary oligonucleotides were annealed and cloned into the BsmBI site of pLentiCRISPRv2 according to established protocols. Exon 5 was selected because it encodes amino acids located between the signal peptide and pore-forming domains of APOL1. Targeting this region was expected to preserve GFP labeling following potential proteolytic processing while minimizing disruption of key APOL1 functional domains.

**Generation and validation of APOL1-edited podocytes**

HIVAN podocytes were seeded in 6-well plates and transfected with 1 µg lentiCRISPRv2 and, where indicated, 1 µg donor vector using Lipofectamine 3000. Twenty-four hours after transfection, cells were selected using 1 µg ml−1 puromycin. Single-cell clones were isolated by limiting dilution. GFP-positive clones were identified and cells were imaged with a Nikon Eclipse TE300 fluorescent microscope, manually isolated, and expanded. Genomic DNA was extracted using DNAzol. Correct GFP integration was confirmed by PCR using the following primer pairs and confirmed by Sanger sequencing:

Left junction:
Forward, 5′-CACCAGTGAATTCACCAAGATGAAGAG-3′
Reverse, 5′-GCCGTCCACGCAGCCCTCCA-3′

Right junction:
Forward, 5′-ACCCGCGAGGACCGCAGCGACGCC-3′
Reverse, 5′-GGCTTCCACCTGGAATCAAC-3′

**APOL1 knockout clones were screened using primers:**

Forward, 5′-TTTCCTTGGTGTGGGAGTGA-3′
Reverse, 5′-CACCCTCCATTCTAAGTGCGA-3′

PCR products were analyzed by AclI digestion and confirmed by Sanger sequencing.

**Construction of GFP-Tagged and Organelle Marker Plasmids.**

For homology-directed repair, ~650–680 bp homology arms flanking APOL1 exon 5 were PCR-amplified from genomic DNA of HIVAN podocyte cells and cloned into the XhoI and SalI sites of pGL3-Basic. GFP coding sequence derived from pZsGreen1-C1 was inserted between the homology arms to generate an in-frame APOL1–GFP donor construct for genome editing.

To generate pCMV-GFP-APOL1-G1, APOL1-G1 cDNA was amplified using the following primers: Forward, 5′-TACGAATTCCATGGAGGGAGCTGCTTTGC-3′; Reverse, 5′-CGCGGATCCTCACAGTTCTTGGTCCGCCTGC-3′. The PCR product was cloned into EcoRI/BamHI-digested pZsGreen1-C1. APOL1-G1 truncation constructs encoding amino acie 1–123 and amino acid 124–398 were generated by the same strategy using the following primer pairs: amino acid 1–123, Forward 5′-TACGAATTCCATGGAGGGAGCTGCTTTGC-3′ and Reverse 5′-TGCAGGTACCTAGATCATTTGTCTTGCAAGGTTGTCC-3′; amino acid 124–398,

Forward 5′-CTTGAATTCCATGAAAGACAAGAACTGGCACG-3′ and Reverse 5′-CGCGGATCCTCACAGTTCTTGGTCCGCCTGC-3′. The APOL1-G1 amino acid 124–398ΔBH3 mutant was generated by overlap-extension PCR using mutagenic primers (Forward, 5′-AGCTTGAGGATAACATAAGAAGGAAGGTCCACAAAGGCACCACCATC-3′; Reverse, 5′-GATGGTGGTGCCTTTGTGGACCTTCCTTCTTATGTTATCCTCAAGCT-3′) together with the amino acid 124–398 amplification primers. For pCMV-GFP-APOL1-G1-NLS, a PCR fragment encoding the APOL1 C terminus fused to the Sam68 nuclear localization sequence (RPSLKAPPARPVKGAYREHPYGRY) was inserted into the PstI/BamHI sites of pCMV-GFP-APOL1-G1.

The ER marker plasmid pRFP-KDEL was generated by replacing GFP with a DsRed-derived coding sequence amplified from pDsRed-Monomer-C1 using NheI and HindIII restriction sites. The lipid droplet marker pRFP-ADRP was generated by replacing GFP in pEGFP-ADRP with DsRed2 using NheI and XhoI restriction sites. All plasmid constructs were confirmed by Sanger sequencing. Complete plasmid maps, annotated sequences and cloning information are available from the corresponding author upon reasonable request.

**Cell proliferation assay**

HIVAN podocytes ($2 \times 10^5$ cells per well) were seeded in 6-well plates. Cells were harvested daily for six days and fixed with 4% paraformaldehyde. Known numbers of red fluorescent reference cells were added before flow cytometric analysis. Absolute cell numbers were calculated from the ratio of podocytes to reference cells.

**RNA sequencing library preparation and sequencing**

RNA sequencing was performed by GENEWIZ (South Plainfield, NJ, USA). Total RNA concentration was determined using the Qubit 2.0 Fluorometer (Thermo Fisher Scientific, Waltham, MA, USA), and RNA integrity was evaluated using the Agilent 4200 TapeStation (Agilent Technologies, Santa Clara, CA, USA). Prior to library preparation, RNA samples were treated with TURBO DNase (Thermo Fisher Scientific) to eliminate contaminating genomic DNA.

Ribosomal RNA depletion was performed using the QIAGEN FastSelect rRNA HMR Kit (Qiagen, Hilden, Germany). Sequencing libraries were generated using the NEBNext Ultra II RNA Library Prep Kit for Illumina (New England Biolabs, Ipswich, MA, USA) following the manufacturer's protocol. Briefly, rRNA-depleted RNA was fragmented at 94 °C for 15 min, followed by first- and second-strand cDNA synthesis. Double-stranded cDNA underwent end repair, 3′ adenylation, adapter ligation, index incorporation, and library enrichment by limited-cycle PCR.

Completed libraries were assessed for fragment size distribution using the Agilent 4200 TapeStation and quantified using both the Qubit 2.0 Fluorometer and quantitative PCR (KAPA Biosystems, Wilmington, MA, USA). Libraries were multiplexed, clustered on an Illumina flow cell, and sequenced on an Illumina HiSeq 3000/4000 platform to generate 2 × 150-bp paired-end reads.

Image processing and base calling were performed using Illumina HiSeq Control Software (HCS). Raw base call (BCL) files were converted to demultiplexed FASTQ files using Illumina bcl2fastq v2.17, allowing one mismatch during sample index assignment.[9]

**Analysis of GFP donor integration**

Potential off-target integration of the GFP donor sequence was assessed using whole-genome sequencing. A custom reference genome was generated by appending the GFP sequence to the human reference genome as an independent contig ("GFP"), and paired-end reads were aligned using BWA-MEM version 0.7.17-r1198-dirty. Read pairs in which one mate aligned to the GFP sequence were extracted and examined for discordant mapping. The intended GFP integration site was located within exon 5 of the APOL1 locus on chromosome 22. No read pairs were identified in which one mate aligned to GFP and the corresponding mate aligned to another chromosome or mapped more than 1 kb from the intended APOL1 insertion site, providing no evidence of detectable off-target GFP integration.

**RNA-sequencing data analysis**

RNA sequencing was performed using four independent biological replicates per group, unless indicated otherwise, generating 42–49 million paired-end reads per sample. Raw sequencing reads were evaluated using FastQC, and quality-control reports were summarized with MultiQC. Adapter sequences were removed using Cutadapt (v3.5) before alignment to the human reference transcriptome (GENCODE release 27) using the splice-aware aligner STAR (version 2.7.11b). [10] Gene-level differential expression analysis was performed in R using DESeq2 (version 1.44.0).[11] following exclusion of low-abundance genes (less than 5 reads in 50% of the samples). Library-size normalization, dispersion estimation, and negative binomial model fitting were performed using the DESeq2 workflow. Variance-stabilized expression values from the 500 most variable genes were used for unsupervised principal component analysis. Differentially expressed genes were ranked by log2 fold change and Benjamini-Hochberg false discovery rate-adjusted *P* values, without excluding any genes based on FDR cutoff, for Gene Set Enrichment Analysis (GSEA). MSigDB version 2026.1.Hs was used as reference pathway database. Pathway enrichment was evaluated using normalized enrichment scores and false discovery rates (FDR <=0.05).

**RNA sequencing data analysis**

RNA sequencing was performed using four independent biological replicates per group unless indicated otherwise, generating 42–49 million paired-end reads per sample. Raw sequencing reads were evaluated using FastQC, and quality-control reports were summarized with MultiQC. Adapter sequences were removed using Cutadapt (version 3.5) before alignment to the human reference transcriptome (GENCODE release 27) using the splice-aware aligner STAR (version 2.7.11b). RNA-Seq reads were aligned to the human reference genome (GRCh38, annotation release 27). Splice junction annotations were provided using an overhang value of 149 bp (--sjdbOverhang 149). Multi-mapping alignments were restricted to a maximum of 20 genomic loci (--outFilterMultimapNmax 20), and alignments with scores within 1 of the top score were retained (--outFilterMultimapScoreRange 1). A maximum of 10 mismatches per read was permitted (--outFilterMismatchNmax 10). The minimum mapped length and score thresholds were set to 33% of the read length (--outFilterMatchNminOverLread 0.33, --outFilterScoreMinOverLread 0.33). Maximum intron and mate gap sizes were capped at 500,000 bp and 1,000,000 bp, respectively. Unmapped reads were retained in the resulting coordinate-sorted BAM files, and raw gene-level counts were mapped simultaneously (--quantMode TranscriptomeSAM GeneCounts). Gene-level differential expression analysis was performed in R using DESeq2 (version 1.44.0)[11] following exclusion of low-abundance genes (less than 5 reads in 50% of the samples). Library-size normalization, dispersion estimation, and negative binomial model fitting were performed using the DESeq2 workflow. Variance-stabilized expression values from the 500 most variable genes were used for unsupervised principal component analysis. Differentially expressed genes were ranked by log2 fold change and Benjamini–Hochberg false discovery rate-adjusted *P* values, without excluding any genes based on FDR cutoff, for Gene Set Enrichment Analysis (GSEA). MSigDB version 2026.1.Hs was used as reference pathway database. Pathway enrichment was evaluated using normalized enrichment scores and false discovery rates (FDR <=0.05).

**Flow cytometry**

For IFN-γ experiments, podocytes were treated with 10 ng $ml^{-1}$ recombinant human IFN-γ or 2 μM inaxaplin (VX-147, Vertex Pharmaceuticals, Boston, MA) for 24 h. Cells were detached using trypsin-EDTA and stained with 5 μM DAPI to identify non-viable cells. For transient transfections, $5 \times 10^5$ cells were seeded per well in 6-well plates and transfected with 3 μg plasmid DNA and 3 μl Lipofectamine 3000.

Flow cytometric analysis was performed on a CytoFLEX Flow Cytometer (Beckman Coulter, Brea, CA) equipped with 405, 488, and 561 nm lasers. Fluorescence of GFP, red and DAPI staining was detected with 525/40 nm, 585/42 nm and 450/50 nm emission filters. A gate was created on FITC channel vs Forward Scatter (FSC) plot to isolate the population of GFP positive cells. The GFP expression was quantified by the Median Fluorescence Intensity (MFI). Dead cells were quantified by gating PB450 channel vs Side Scatter (SSC) plot for population of DAPI stained cells in GFP positive cells. Debris and doublets were excluded before analysis, and GFP positive cells were gated prior to viability analysis. The gates were held constantly for all samples. A minimum of $3 \times 10^5$ events were collected and data were analyzed using FlowJo v10.6.2 software (BD Biosciences, Milpitas, CA).

**Genomic DNA isolation**

High-molecular-weight genomic DNA was isolated from cultured cells using a standard phenol–chloroform extraction protocol. Briefly, cell pellets were lysed in proteinase K-containing lysis buffer, followed by phenol–chloroform extraction and ethanol precipitation. DNA was resuspended in nuclease-free water, and concentration and purity were determined by Qubit fluorometry (Thermo Fisher Scientific) and spectrophotometric analysis before library preparation.

**Whole-genome sequencing library preparation**

Genomic DNA quality control, whole-genome library preparation, and sequencing were performed by Novogene (Sacramento, CA, USA). DNA concentration was measured using a Qubit fluorometer, while DNA integrity and purity were assessed by agarose gel electrophoresis.

Whole-genome sequencing libraries with an average insert size of approximately 350 bp were prepared using the NEBNext Ultra II DNA Library Prep Kit for Illumina (New England Biolabs, Ipswich, MA, USA). Briefly, genomic DNA was randomly fragmented, followed by end repair, A-tailing, Illumina adapter ligation, PCR amplification, size selection, and purification. Library quality was evaluated using Qubit fluorometry for DNA quantification, quantitative PCR for library quantification, and an Agilent Bioanalyzer for fragment size distribution.

Qualified libraries were pooled and sequenced on an Illumina NovaSeq platform using 2 × 150-bp paired-end (PE150) chemistry to achieve an average genome coverage of approximately 40×. Following sequencing, raw reads were subjected to quality filtering by removing reads containing adapter sequences, reads with more than 10% ambiguous bases (N), and reads in which more than 50% of bases had a Phred quality score ≤5. The resulting high-quality sequencing reads were retained in FASTQ format for downstream analyses. We used the human reference genome (GRCh37) with decoy [human_g1k_v37_decoy.fasta] [Data set]. Retrieved from ftp://ftp.broadinstitute.org/bundle/b37/)

**Sequence alignment and variant analysis**

Whole-genome sequencing generated a mean genome coverage of 31–57×, with more than 500 million paired-end 150-bp reads obtained for each sample. Raw sequencing reads were evaluated using FastQC to assess per-base sequence quality, GC content, sequence duplication levels, and adapter contamination. High-quality reads were aligned to the human reference genome hs37d5 (GRCh37) with decoy sequences (*human_g1k_v37_decoy.fasta*) using BWA-MEM (v0.7.17-r1198-dirty). PCR duplicates were identified and marked using Picard MarkDuplicates (v2.27.5-1-gcfb8749-SNAPSHOT).[10]

Single-nucleotide variants (SNVs) and small insertions/deletions (indels) were identified using the Genome Analysis Toolkit (GATK) HaplotypeCaller joint genotyping workflow following the GATK Best Practices recommendations (GATK v4.6.2.0).[12] Base Quality Score Recalibration (BQSR) was performed using dbSNP build 138 (*dbsnp_138.b37.vcf*) as the known-sites resource to correct systematic sequencing errors. Variant Quality Score Recalibration (VQSR) was subsequently applied to distinguish high-confidence variants from technical artifacts. Variants passing quality-control filters were annotated using Ensembl Variant Effect Predictor (VEP) (release 106 for GRCh37),[12] which assigned predicted functional consequences and dbSNP reference SNP identifiers (rsIDs). Annotated variants were further analyzed using GEMINI (0.30.2)[13] for downstream filtering, prioritization, and interpretation.

Per-sample sequencing quality metrics, including sequencing depth, genome coverage, alignment statistics, duplicate read rates, and variant-calling metrics, are summarized in

Supplementary Table S1. Whole-genome sequencing data were additionally used to assess potential off-target integration of the GFP donor cassette, with results presented in Supplementary Fig. 2.

**Protein C-terminal sequencing by liquid chromatography-mass spectrometry (LC-MS/MS)**

Nuclear extracts were prepared using the Abcam Cell Nuclear Extraction Kit (#ab219177). APOL1-containing protein complexes were immunoprecipitated using Proteintech anti-APOL1 antibody and Dynabeads Protein A (#10006D; Thermo Fisher Scientific).

Samples were separated by SDS–PAGE and stained with Coomassie Brilliant Blue. Bands corresponding to GFP–APOL1 were excised and analyzed by MtoZ Biolabs (Boston, MA) using in-gel digestion followed by nano LC–MS/MS analysis. The GFP-APOL1 protein band was excised from SDS–PAGE gels, cut into approximately 1 $mm^3$ pieces, and repeatedly destained using 50% acetonitrile in 50 mM ammonium bicarbonate until colorless. Gel pieces were dehydrated with 100% acetonitrile before reduction with 10 mM dithiothreitol for 1 h at 56°C and alkylation with 55 mM iodoacetamide for 1 h at room temperature in the dark. Following dehydration, gel pieces were digested overnight (16 h at 37°C) using either sequencing-grade trypsin (25 ng $\mu L^{-1}$) or sequencing-grade chymotrypsin (500 ng $\mu L^{-1}$). Peptides were extracted twice using 5% trifluoroacetic acid, 50% acetonitrile, and 45% water, followed by vacuum centrifugation and desalting using self-packed reversed-phase desalting columns. NanoLC separation was performed on a Vanquish Neo ultra-high-performance liquid chromatography system (Thermo Fisher Scientific) equipped with a 100 µm × 180 mm C18 analytical column packed with ReproSil-Pur 120 C18-AQ (3 µm). Peptides were separated at a flow rate of 600 nL $min^{-1}$ using solvent A (0.1% formic acid in water) and solvent B (0.1% formic acid in 80% acetonitrile). The gradient increased from 4% to 40% solvent B over 120 min before column washing at 95% solvent B.

Mass spectrometry was performed using an Orbitrap Fusion Lumos Tribrid mass spectrometer operating in positive-ion, data-dependent acquisition mode. Full MS scans were acquired at 120,000 resolutions over an m/z range of 300–1800. Precursors were fragmented by higher-energy collisional dissociation (HCD; normalized collision energy 30%), and MS/MS spectra were acquired in the Orbitrap at 15,000 resolutions. Raw data were processed using Byonic software (Protein Metrics). Searches were performed using both tryptic and chymotryptic digestion specificity with up to three missed cleavages. Carbamidomethylation of cysteine was designated as a fixed modification, whereas methionine oxidation and protein N-terminal acetylation were included as variable modifications. Precursor and fragment mass tolerances were set to 20 ppm and 0.02 Da, respectively. The C-terminal peptide spectrum was manually reviewed to verify sequence assignment. The identified terminal peptide (ALDNLARQMI) demonstrated that the recombinant GFP–APOL1 construct terminated at amino acid residue 404.

**Cathepsin S cleavage assay**

Recombinant APOL1 (2.5 μg.) was incubated with recombinant cathepsin S 0.16 μg in a 20 μl reaction containing 250 mM MES, 100 mM DTT and 50 mM EDTA (pH 6.0) at 37 °C for 1 h in the presence or absence of 1 mM cathepsin S inhibitor (Apexbio Technology #1373215-15-6). Reaction products were analyzed by SDS–PAGE followed by Coomassie staining or immunoblotting using APOL1 monoclonal antibodies 3.1C1 A80C /3.7D6 P80C (Genentech).

**Statistics**

Data are presented as mean ± s.d. Statistical analyses were performed using GraphPad Prism (v11). Comparisons between two groups were performed using unpaired two-tailed Welch's *t*-tests. Comparisons among three or more groups were performed using one-way analysis of variance (ANOVA) followed by Tukey's multiple-comparison test. All statistical tests were two-sided unless otherwise indicated. $P < 0.05$ was considered statistically significant. For RNA-sequencing analyses we used Benjamini–Hochberg adjustment and GSEA false-discovery rates. The number of biological replicates (*n*), statistical tests, and exact *P* values are provided in the corresponding figure legends.

**References.**


1 Soler-Garcia, A. A., Johnson, D., Hathout, Y. & Ray, P. E. Iron-related proteins: candidate urine biomarkers in childhood HIV-associated renal diseases. *Clin J Am Soc Nephrol* **4**, 763–771 (2009). https://doi.org/10.2215/CJN.0200608

2 Beng, H. *et al.* HIV-Associated CKDs in Children and Adolescents. *Kidney Int Rep* **5**, 2292–2300 (2020). https://doi.org/10.1016/j.ekir.2020.09.001

3 Xie, X. *et al.* The basic domain of HIV-tat transactivating protein is essential for its targeting to lipid rafts and regulating fibroblast growth factor-2 signaling in podocytes isolated from children with HIV-1-associated nephropathy. *J Am Soc Nephrol* **25**, 1800–1813 (2014). https://doi.org/10.1681/ASN.2013070710

4 Ray, P. E. *et al.* Infection of human primary renal epithelial cells with HIV-1 from children with HIV-associated nephropathy. *Kidney Int* **53**, 1217–1229 (1998). https://doi.org/10.1046/j.1523-1755.1998.00900.x

5 Li, J. *et al.* Transmembrane TNF-alpha Facilitates HIV-1 Infection of Podocytes Cultured from Children with HIV-Associated Nephropathy. *J Am Soc Nephrol* **28**, 862–875 (2017). https://doi.org/10.1681/ASN.2016050564

6 Cong, L. *et al.* Multiplex genome engineering using CRISPR/Cas systems. *Science* **339**, 819–823 (2013). https://doi.org/10.1126/science.1231143

7 Mali, P. *et al.* RNA-guided human genome engineering via Cas9. *Science* **339**, 823–826 (2013). https://doi.org/10.1126/science.1232033

8 Scales, S. J. *et al.* Apolipoprotein L1-Specific Antibodies Detect Endogenous APOL1 inside the Endoplasmic Reticulum and on the Plasma Membrane of Podocytes. *J Am Soc Nephrol* **31**, 2044–2064 (2020). https://doi.org/10.1681/ASN.2019080829

9 Van der Auwera, G. A. *et al.* From FastQ data to high confidence variant calls: the Genome Analysis Toolkit best practices pipeline. *Curr Protoc Bioinformatics* **43**, 11 10 11–11 10 33 (2013). https://doi.org/10.1002/0471250953.bi1110s43

10 Dobin, A. *et al.* STAR: ultrafast universal RNA-seq aligner. *Bioinformatics* **29**, 15–21 (2013). https://doi.org/10.1093/bioinformatics/bts635

11 Love, M. I., Huber, W. & Anders, S. Moderated estimation of fold change and dispersion for RNA-seq data with DESeq2. *Genome Biol* **15**, 550 (2014). https://doi.org/10.1186/s13059-014-0550-8

12 McLaren, W. *et al.* The Ensembl Variant Effect Predictor. *Genome Biol* **17**, 122 (2016). https://doi.org/10.1186/s13059-016-0974-4

13 Paila, U., Chapman, B. A., Kirchner, R. & Quinlan, A. R. GEMINI: integrative exploration of genetic variation and genome annotations. *PLoS Comput Biol* **9**, e1003153 (2013). https://doi.org/10.1371/journal.pcbi.1003153
